# An ELISA-based method for rapid genetic screens in *Drosophila*

**DOI:** 10.1101/2021.04.21.440847

**Authors:** Taylor R Jay, Yunsik Kang, Amanda Jefferson, Marc R Freeman

## Abstract

Drosophila is a powerful model in which to perform genetic screens, but screening assays that are both rapid and can be used to examine a wide variety of cellular and molecular pathways are limited. *Drosophila* offer an extensive toolbox of GFP-based transcriptional reporters, GFP-tagged proteins, and driver lines which can be used to express GFP in numerous subpopulations of cells. Thus, a tool that can rapidly and quantitatively evaluate GFP levels in *Drosophila* tissue would provide a broadly applicable screening platform. To quantify GFP levels from *Drosophila* lysates, we developed a GFP-based ELISA assay. We demonstrate that this assay can detect membrane localized GFP in a variety of neuronal and glial cell populations and validate that it can identify genes that change the morphology of these cells. This assay was also able to detect STAT transcriptional activity after injury. We found that this assay can detect endogenously GFP-tagged proteins, including Draper and Cryptochrome, and it is able to report developmental and circadian changes in the expression of these proteins. Finally, we validated that the assay can be used to detect changes in synapse elimination upon genetic manipulation of astrocytes. We then used the assay to perform a small-scale screen, which identified Syntaxins as novel regulators of astrocyte-mediated synapse elimination. Together, these studies establish an ELISA as a rapid, easy and quantitative *in vivo* screening method to assay a wide breadth of fundamental questions in neurobiology.

**Significance Statement:** Forward genetic screens in *Drosophila* have played an integral role in elucidating the cellular and molecular pathways that govern almost every facet of biology. However, current screening methods in *Drosophila* are either fast, but limited in their specificity for particular pathways or processes, or rely on imaging, which requires substantial expertise, time, and cost. We have developed a rapid GFP-based ELISA screening method that, when paired with the wealth of GFP-based genetic tools already available in *Drosophila*, can be used to screen for regulators of many subpopulations of cells, transcriptional programs and levels of thousands of different proteins. Using this assay, we have identified a novel family of genes required for astrocytes to mediate developmental synapse elimination. This technique provides a screening platform that is fast, accessible, and broadly applicable to many pathways and processes, making *Drosophila* an even more powerful screening tool.

## Introduction

*Drosophila* are an excellent model system for forward genetic screens. Their fast generation time, balance of relative genetic simplicity and cellular and organismal complexity, and the high degree of conservation in genes and pathways between *Drosophila* and vertebrates, make *Drosophila* a powerful system for gene discovery.

Historically, there has been a tradeoff between designing screens with efficient read outs, and designing screens to identify genes specifically involved in a desired phenotype of interest (1). In the first category, many classic and extremely productive screens have been performed using easy to identify phenotypes, such as lethality (2) or sterility (3). Others devised behavioral assays that could be performed with many flies in parallel (4-6), or studied processes that affect easily recognizable external features, such as using patterning of the fly’s cuticle to screen for genes involved in embryonic development (7) or studying genes that are involved in gross development of the eye (8). These types of screens can be performed rapidly, but are limited in the specificity of the cellular and molecular pathways of the genes they identify.

In order to interrogate more specific questions, screens in *Drosophila* have traditionally relied on imaging based assays. These can be performed in a variety of different tissues and cell types and assess a wide variety of molecular pathways (9, 10). In these screens, the tissue of interest is often dissected, labeled, and then samples are imaged and are typically qualitatively scored by the investigator. Even in systems designed to dramatically reduce the time required for these screens (11), they still take multiple person-years to complete. These screens are also costly and require specialized skills.

To address the limitations of these current approaches, we adapted an enzyme-linked immunosorbent assay (ELISA) to work with *Drosophila* lysates. ELISAs provide a quantitative read out of the concentration of a protein of interest, and thus can be used to answer a wide variety of specific molecular questions. ELISAs are fast and high throughput, require little technical expertise, and provide quantitative results. In addition, compared to imaging-based screens, they are relatively inexpensive and require no specialized equipment, making this screening method accessible to a wide variety of laboratory environments.

ELISAs have been used extensively for rapid screening *in vitro*. However, many questions simply cannot be addressed in cultured cells. Cells *in vitro* often lack the complex morphology observed *in vivo*, and thus cannot be used to investigate how diverse cell types establish and maintain these morphological features. Many cells also undergo large-scale transcriptional changes when cultured, making it difficult to study regulation of their transcriptional programs. Even in co-culture systems, it is challenging to assess questions about how different cell types interact in complex tissues *in vitro*. Finally, it is difficult to accurately recapitulate complex organismal events, like development, injury or disease in a dish. *Drosophila* provides a highly genetically tractable system in which these questions can be studied *in vivo*, and so adapting an ELISA assay for use with *Drosophila* lysates extends the types of questions that can be addressed using this rapid screening method.

While ELISAs can be used to identify many protein targets, we chose to develop an assay to quantify levels of GFP to maximize its versatility. Paired with the large array of GFP-based tools already available in *Drosophila*, this assay can be used to address a wide variety of cellular and molecular questions with minimal or no troubleshooting of new assay conditions. GFP can be driven in most cellular compartments and in almost any desired cell type using the vast array of Gal4 and other driver lines that have been developed (12-16), allowing a GFP-based ELISA to potentially assess the abundance of virtually any subpopulation of cells and their subcellular components. There are GFP-based transcriptional reporters that can be used to assess activity of a variety of different transcription factors, as well as reporters of signaling pathways and cellular metabolism and physiology (17-20). In addition, multiple approaches have been used in ongoing, large-scale efforts to insert endogenous GFP tags into all conserved *Drosophila* proteins (21-25). Thus, a GFP-based ELISA assay could be used to screen for regulators of almost any protein in any tissue type.

Here, we provide six independent examples of how this assay can be used to assess cellular morphology, transcriptional regulation, and protein levels in the *Drosophila* nervous system. Further, we apply this assay to perform a small-scale screen that reveals new biology about interactions between astrocytes and neurons during the process of developmental synapse elimination.

## Results

### Validation of a GFP-Based ELISA Assay for Use with *Drosophila* Tissue

In order to use an ELISA as a screening method, we first needed to optimize and validate its use with *Drosophila* lysates. After optimizing tissue preparation and assay conditions, we were able to achieve substantial enrichment of signal in lysates prepared from homogenized heads of flies expressing GFP in all neurons (Fig 1A), relative to lysates prepared from wild type fly heads (Fig 1B, Fig S1A), even with relatively low antibody concentrations. To evaluate the detection limits of the method, we used recombinant GFP to assay its dynamic range (Fig 1C) and established that the assay was sufficiently sensitive to detect concentrations as low as 25 pg/mL GFP (Fig S1B). Using *Drosophila* tissue, we found that the assay could detect signal ranging from 1-10 heads of flies expressing GFP in all neurons within the assay’s linear dynamic range (Fig 1D). We developed this in-house ELISA assay using commercially available antibodies and solutions that are commonly available in most laboratories to ensure it was cost-effective to perform large-scale screens. However, we found that a rapid commercial GFP ELISA kit can also perform well at detecting GFP in *Drosophila* lysates (Fig S1C). While the cost of this commercial assay is approximately 25 times higher than our assay, it can be performed in under two hours, compared to our assay which uses a multi-day protocol. Together, these studies demonstrate that an ELISA assay can robustly detect GFP signal in *Drosophila* tissue, with high resolution between GFP expressing flies and non-GFP expressing controls, even in single fly heads. This assay also has a linear dynamic range that allows signal detection across a sufficient scale to detect physiologically meaningful differences in GFP levels.

**Figure 1.**
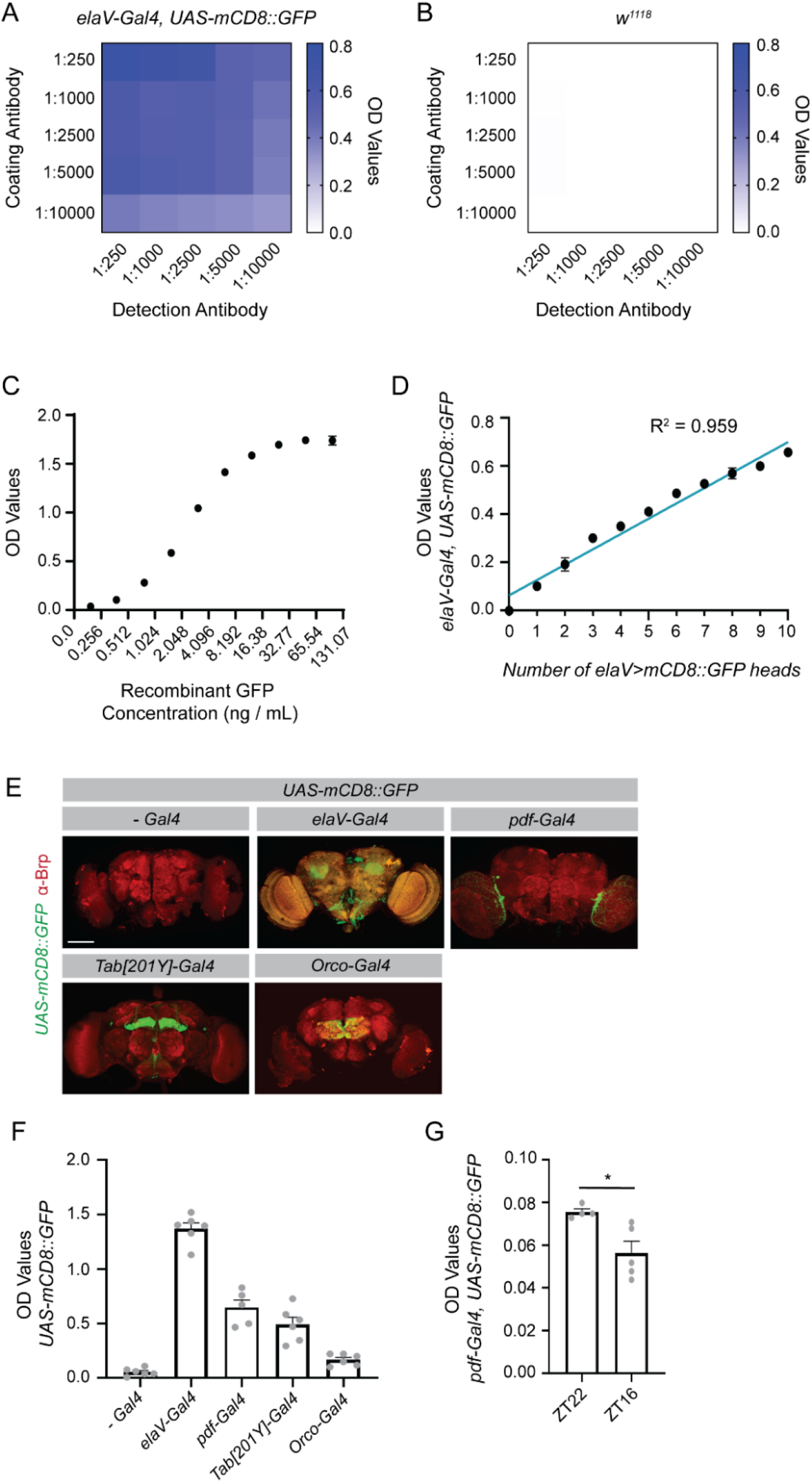
An ELISA assay can be used to quantify GFP levels in Drosophila in diverse cell populations. An ELISA assay was optimized to detect GFP in Drosophila lysates. **(A)** For each sample, lysates were prepared from 5 heads collected from flies expressing a membrane localized GFP under the control of the pan-neuronal *elaV* driver. Optical density (OD) values were recorded with different concentrations of coating antibody (chicken α GFP) and detection antibody (mouse α GFP). Depicted OD values represent the mean of two biological replicates. **(B)** A parallel analysis was performed as in (A) but in lysates containing 5 heads from *w*^*1118*^ flies. **(C)** Indicated concentrations of recombinant GFP were evaluated by ELISA and the resulting OD values are shown. Six replicates were performed for each concentration. **(D)** Ten fly heads were prepared for each sample, with the number on the x-axis representing the number of heads from *elaV>mCD8::GFP* expressing flies. The remaining heads in each sample were from *w*^*1118*^ flies. OD values were recorded from 3 replicates. A correlation analysis was performed using a simple linear regression. **(E)** Flies expressing the indicated drivers were crossed with *UAS-mCD8::GFP* flies, their brains dissected at 10-12 dpe and stained for Brp. The GFP depicted is from the endogenous GFP signal. Images were acquired at 20x magnification and represent maximum projections of z slices taken every 3μm through the brain. **(F)** Five fly heads from each of the indicated genotypes were prepared for each sample and the OD values assessed by ELISA. **(G)** Five fly heads were collected for each sample from flies expressing a membrane localized GFP under the control of the *pdf* driver. Flies were collected at zeitgeber time (ZT) 22, shortly before the onset of light, and at ZT16 shortly after the onset of the dark phase. OD values for these samples were assessed by ELISA. Graphs represent the mean ± SEM. *p<0.05. Scale bar - 100μm.

### Detecting GFP Expression in Small Subpopulations of Cells and Correlates with Circadian Changes in Neuronal Morphology

While we had established that the ELISA assay could be used to detect GFP when expressed in all neurons, we next wanted to assess whether the assay was also sensitive enough to detect GFP when expressed in smaller subpopulations of cells. We homogenized heads from flies expressing a membrane localized GFP under the control of several Gal4 drivers that express in subpopulations of neurons. GFP expression was readily detectable in all populations tested, including mushroom body neurons, olfactory receptor neurons, and even the approximately 150 *pdf* expressing neurons (Fig 1E, F). We next wanted to test whether this assay could be used to detect relevant changes in these neuronal subpopulations. It has previously been shown that *pdf* neurons undergo circadian remodeling, with elimination of a subset of their processes in the early dark phase (26). To determine whether we could detect these circadian differences using the ELISA assay, we collected heads from flies at zeitgeber time (ZT) 22, just preceding the onset of the light phase and ZT16, early in the dark phase. We were able to detect a significant decrease in GFP expression between these two time points (Fig 1G), in line with the imaging studies that established a reduction in processes across circadian time. Thus, this assay could be used to screen for molecules that regulate circadian remodeling of these neurons. The signal detection in the additional driver lines used here also suggest that the assay is likely to be compatible with many of the thousands of cell-specific driver lines available in *Drosophila*, which would allow one to explore a diverse array of questions in a variety of cell populations using quantification of GFP levels by ELISA as an initial readout.

### Identifying Molecules Involved in Establishing Complex Cellular Morphology

It has long been a challenge to screen for molecules involved in establishing the complex morphology of cells like astrocytes, which fail to recapitulate these morphologies *in vitro*. We wanted to test whether we could use our ELISA assay to identify regulators of astrocyte morphology *in vivo*. Using a MARCM-based clonal expression system, it has previously been established that signaling by ligands Thisbe (Ths) and Pyramus (Pyr) through the FGF receptor Heartless (Htl) promotes astrocyte growth (27). Indeed, when we expressed a membrane-localized GFP in astrocytes, we were able to detect by imaging increased astrocyte area within the neuropil in flies in which astrocytes overexpress *ths* and *pyr* and in flies which express a constitutively active form of the Htl receptor (Fig 2A, B). Conversely, knockdown of *htl* resulted in reduced astrocyte area (Fig 2A, B). These findings were consistent with quantification of images using antibody staining for GAT (Fig S2A, S2B), a membrane-localized astrocyte marker (27), suggesting that the GFP signal reliably correlates with astrocyte membrane area. We were able to recapitulate all of the differences observed in these imaging studies by measuring GFP signal using the ELISA assay (Fig 2C). Not only did this assay require fewer animals and less time, but the variability in these results was much lower than when using image quantification (Fig 2B, C). We did not detect differences in GFP signal by ELISA with different numbers of *UAS* sequences in the background (Fig S2C), indicating that *UAS* dosage effects were not responsible for the differences we observed in the experimental genotypes used in this study. These results establish that the ELISA assay can be used to identify genes that alter astrocyte morphology, and could be used as an initial screening platform to identify novel regulators of astrocyte morphogenesis and maintenance. It could also be applied to screens aimed at identifying genes required to establish the morphology of other cell types in an *in vivo* system.

**Figure 2.**
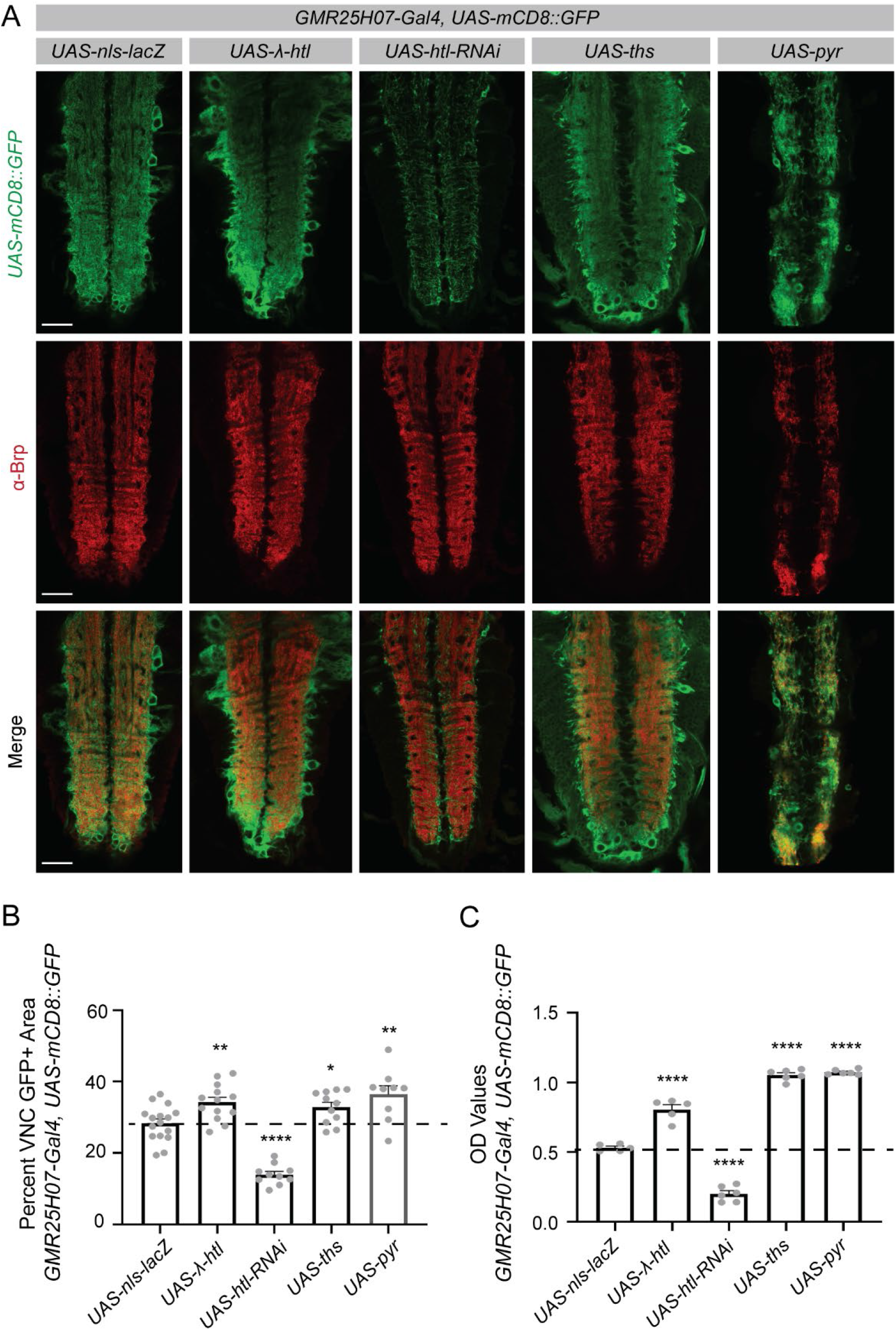
The ELISA assay can be used to screen for changes in complex cellular morphology. The driver line, *GMR25H07-Gal4* in which Gal4 is expressed under the control of an enhancer associated with the astrocytic *alrm* gene, was used to express a membrane localized GFP and to manipulate expression of genes previously implicated in regulating astrocyte morphogenesis. **(A)** Images of the larval ventral nerve cord (VNC) were acquired at 20x magnification and single z planes are shown for each genotype. Immunohistochemistry was performed for GFP to detect the membrane localized reporter, and for Brp to identify the neuropil of the VNC. **(B)** GFP+ area within the VNC neuropil was quantified using single z planes, as shown above. **(C)** The CNS was dissected from 3 larvae for each sample and the GFP signal within these samples quantified by ELISA. Graphs represent mean ± SEM and each experimental genotype was statistically compared to the UAS-nls-lacZ control sample (mean depicted as a dotted line). *p<0.05, **p<0.01, ***p<0.001, ****p<0.0001. Scale bar - 50μm.

### Quantifying Signal from GFP-Based Transcriptional Reporters

There is a large collection of GFP-based transcriptional reporters available in *Drosophila*, and we sought to determine whether we could use the ELISA assay to measure GFP signal from these transcriptional reporter tools. As an example, we chose to use a *STAT92E*-based GFP reporter, in which a destabilized GFP is expressed under the control of *STAT92E* enhancer sequences (19). We found that using this destabilized GFP was necessary to use in this context because the long half-life of GFP resulted in high levels of signal at baseline. This line has previously been used to evaluate upregulation in STAT signaling following injury in *Drosophila* (28). Using imaging, we were able to recapitulate the expected increase in GFP signal within glial cells surrounding severed axons in the antennal lobe following antennal ablation (Fig 3A). While we were able to see this injury-induced increase by imaging, we did need to amplify the signal using a GFP antibody, indicating that this reporter is relatively weakly activated in this context.

**Figure 3.**
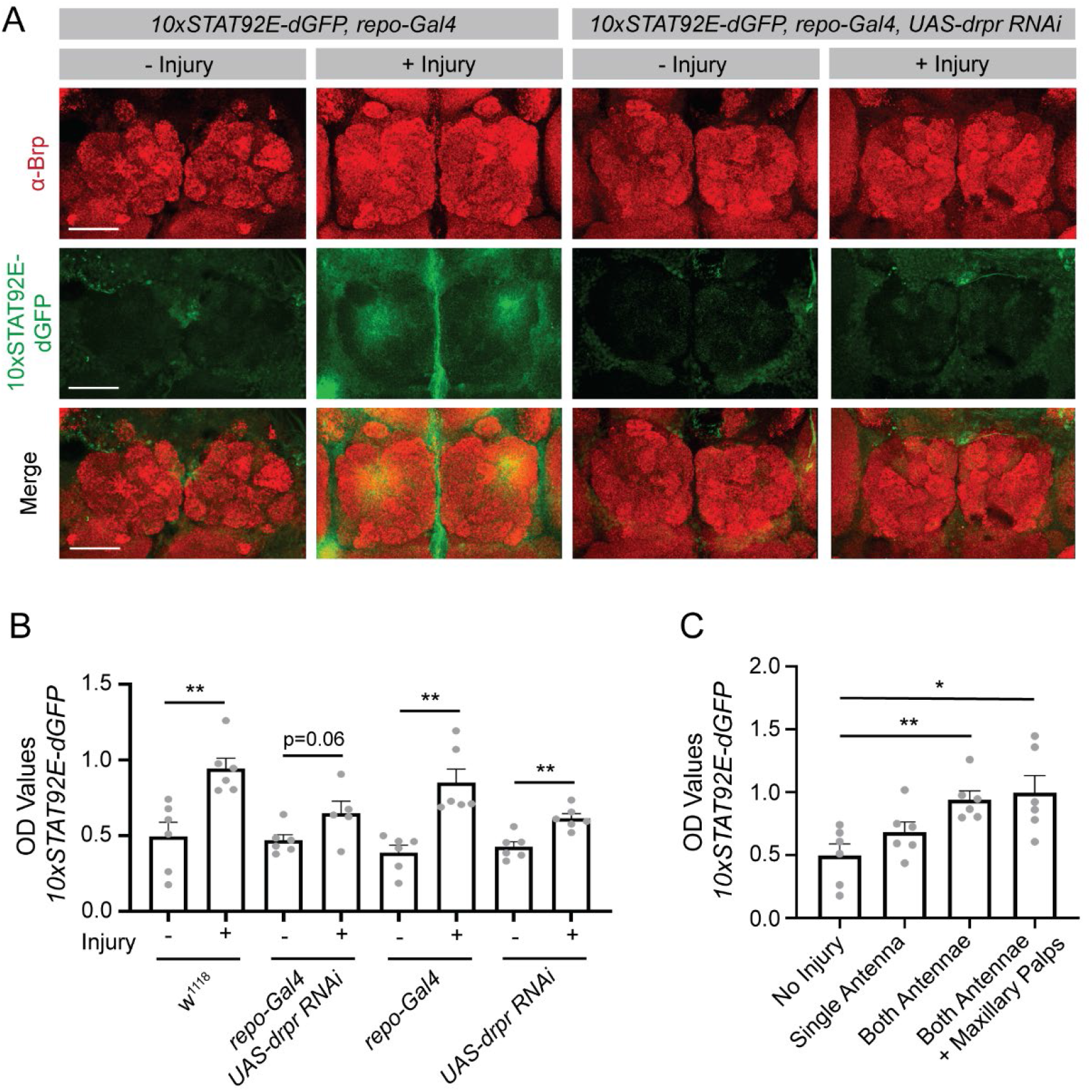
The ELISA assay can be used to screen for transcriptional regulators using GFP transcriptional reporters. The *STAT92E-dGFP* reporter was expressed ubiquitously and gene expression manipulated in all glia using the *repo-Gal4* driver. **(A)** Both antennae were removed from flies at 9-11 dpe and their brains were dissected after 24 ± 2 hours. Brains were stained for Brp to label the neuropil and GFP to identify reporter activity. Images were acquired at 20x and a maximum projection of z sections taken 1μm apart through the antennal lobes are shown. **(B)** Flies were injured as described in A and 3 fly heads were collected for each sample 24 hours after injury. Lysates were prepared, and GFP OD values assessed by ELISA. Statistical comparisons were performed between uninjured and injured flies for each genotype. **(C)** Graded severities of injury, as indicated on the x-axis, were performed in flies expressing the STAT92E-GFP reporter and GFP OD values were measured by ELISA. Statistical comparisons were performed between the no injury condition and each of the injury conditions. Graphs represent mean ± SEM. *p<0.05, **p<0.01, or as indicated. Scale bar - 50μm.

Despite this, we were able to detect robust increases in GFP signal by ELISA after injury, likely due to the high sensitivity of the ELISA assay (Fig 3B). This response was attenuated by expression of an RNAi directed against *draper* (*drpr)* in glia (Fig 3A, B), consistent with Drpr’s known role in mediating glial injury responses (28). We were also able to detect graded responses in signal from the transcriptional reporter by ELISA in response to increasing severities of injury (Fig 3C). These results demonstrate that the ELISA can be used to detect glial transcriptional changes in response to injury in *Drosophila*, and could be used to screen for genes involved in neuronal signaling that an injury has occurred or glial responses to those signals. More broadly, these results demonstrate that our ELISA assay can be used to measure signal from GFP-based transcriptional reporters, even those that are activated at relatively low levels.

### Detecting Proteins with Endogenously Expressed GFP Tags

*Drosophila* have a wealth of endogenously GFP-tagged proteins available, with the eventual goal of tagging every conserved protein with GFP (22-25, 29). Other strategies have also created comprehensive libraries of GFP tagged proteins inserted as BACs (30, 31) or fosmids (32). Thus, if the ELISA assay could detect these endogenously tagged proteins, it could in principle be used to screen for regulators of virtually any protein of interest. We tested whether it could be used to detect expression of two example endogenously GFP tagged proteins, Drpr which was tagged within the endogenous *drpr* locus, and Cryptochome (Cry) which was inserted with a GFP tag using a BAC.

It had previously been shown that the phagocytic protein Drpr is upregulated during metamorphosis (33), a developmental period during which the nervous system undergoes substantial remodeling, thereby placing high phagocytic demands on glia to clear extensive cellular debris. Indeed, we found using imaging that flies exhibited a substantial increase in Drpr-GFP expression during the period of metamorphosis from 2 to 6 hours after puparium formation (APF) (Fig 4A), similar to what we detected using an antibody against Drpr (Fig 4B). This upregulation was attenuated when astrocytes expressed a dominant negative form of the Ecdysone Receptor (EcR^DN^), which largely prevents these cells from responding to the cue that drives metamorphosis. Using the ELISA assay, we were similarly able to detect a robust increase in GFP signal in flies expressing Drpr-GFP from 2 to 6 hours APF, and this response was partially attenuated in flies expressing EcR^DN^ in astrocytes (Fig 4C), similar to what we observed by imaging. This validates that the assay can detect GFP signal from flies expressing endogenously GFP tagged Drpr, and that it is able to report robust increases in protein expression over the course of development. This could be used to screen for genes involved in establishing appropriate regulation of Drpr protein levels, or more broadly for genes that are required for glia to respond to developmental cues to change their function in appropriate contexts.

**Figure 4.**
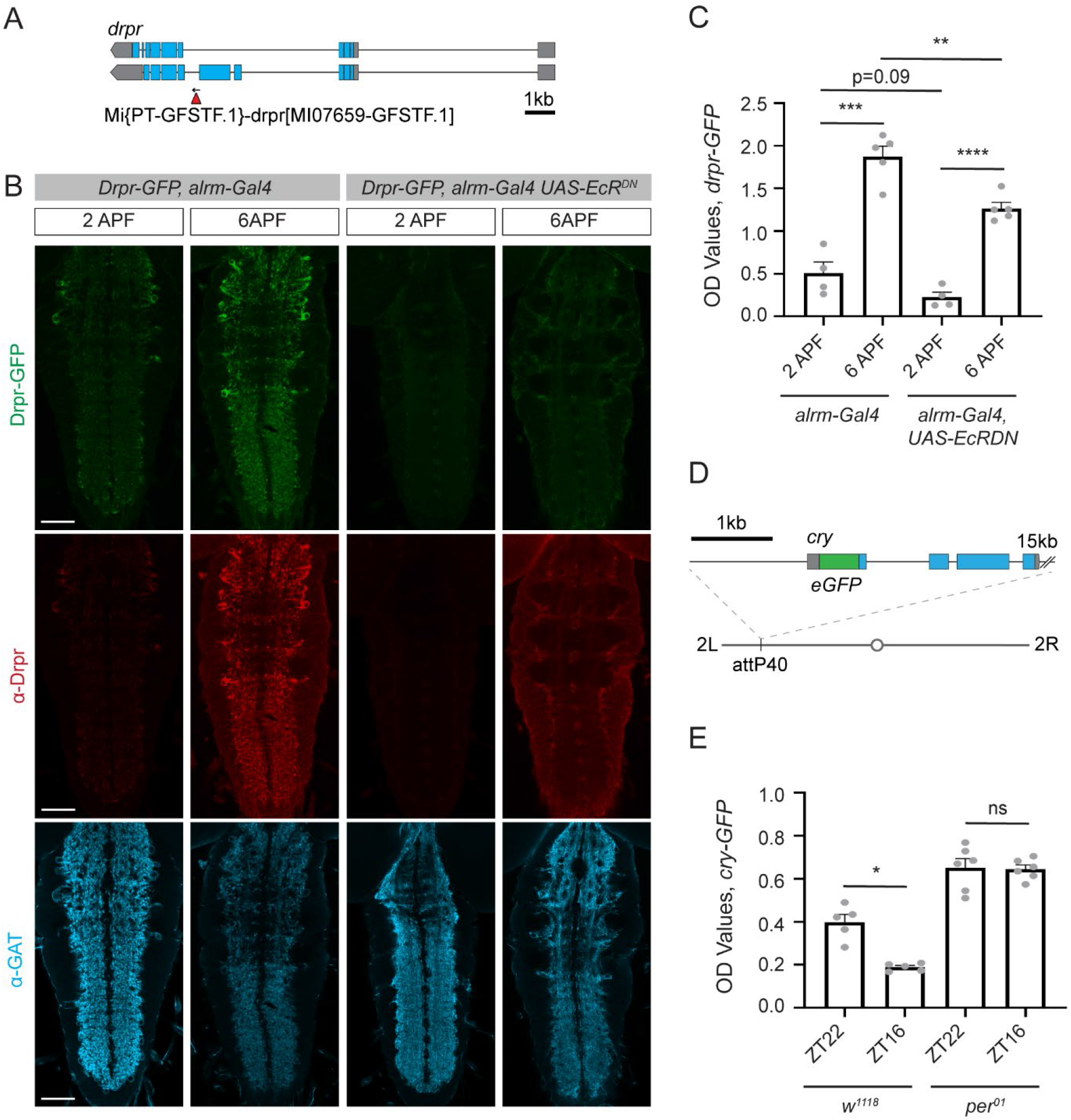
The ELISA assay can be used to detect endogenously GFP tagged proteins. The ELISA was used to detect levels of two endogenously GFP tagged proteins. **(A)** A line expressing Draper-GFP (Drpr-GFP) was previously generated using a MiMIC insertion of GFP into the endogenous *drpr* locus. **(B)** Drpr levels were assessed at 2 hours after puparium formation (2APF) and at 6APF. This was performed in a control background, in which the astrocyte driver *alrm-Gal4* was expressed alone, and in flies expressing a dominant negative ecdysone receptor (*EcR*^*DN*^) under the control of the *alrm-Gal4* driver. Pupae were dissected and immunohistochemistry was performed for Drpr and the astrocytic marker GAT. The GFP signal represents the endogenous signal from Drpr-GFP. Images were acquired at 20x magnification and represent a single z plane of the VNC. **(C)** The CNS from 5 pupae were dissected for each sample and GFP OD levels assessed by ELISA. **(D)** A line expressing Cryptochrome-GFP (Cry-GFP) was previously generated by inserting a GFP sequence into the N-terminus of *cry* in a plasmid containing a 20kb genomic region which includes the *cry* gene. This sequence was then inserted at the attP40 site on the second chromosome. **(E)** Cry-GFP levels were assessed by ELISA from 3 heads per sample collected at zeitgeber time (ZT22) just preceding the onset of light and at ZT16, shortly after the beginning of the dark phase. This was also performed in a *per*^*01*^ mutant background. Graphs represent mean ± SEM. ns-not significant, *p<0.05, **p<0.01, ***p<0.001, ****p<0.0001, or as indicated. Scale bar - 50μm.

We next attempted to use the ELISA to measure a GFP tagged Cry (Fig 4D), which was inserted in a BAC and was previously shown to act as a functional Cry protein (34) and to maintain its endogenous pattern of circadian regulation (35). We were able to detect this endogenously tagged Cry-GFP in *Drosophila* heads using the ELISA, and similar to what had been previously described, found that its levels were higher at circadian times shortly before the onset of light (ZT22) and lower early in the dark period (ZT16) (Fig 4E). Mutants in the gene *period* (*per*) have previously been shown to abrogate circadian rhythms (6), and we show that *per*^*01*^ mutants no longer show circadian changes in Cry protein expression by ELISA (Fig 4E). Our ELISA results also indicate that Cry protein levels are elevated in *per*^*01*^ mutants, consistent with the known role for Cry and Per complexes in inhibiting *cry* transcription (35). These results demonstrate that the ELISA can detect endogenously GFP tagged Cry protein, and report expected circadian changes in its expression. Thus, this assay could be used to screen for modifiers of circadian regulation of Cry protein. Taken together, these results provide a proof of principle that the ELISA assay can be used to detect endogenously GFP tagged proteins, which promises to provide a platform to screen for regulators of many of the thousands of proteins for which GFP tagged versions are available.

### Identifying Novel Classes of Glial Genes Involved in Regulation of Synapse Elimination

We took advantage of the ability of the ELISA to detect endogenously GFP tagged proteins to use it to detect synapses using a GFP tagged Bruchpilot (Brp) (Fig 5A), a protein integral to presynaptic structure (36) which is commonly used to evaluate synapses by immunohistochemistry (37). We specifically wanted to evaluate glial genes involved in the regulation of synaptic engulfment. Work in mammalian systems has established critical roles for glial synaptic engulfment in nervous system development, and in modulating neurological diseases (38-40). However, it is difficult in mammalian systems to simultaneously manipulate neuronal and glial gene expression *in vivo*, and it is extremely time and cost-intensive to perform forward genetic screens in mammals. Thus, our knowledge about how neurons and glia communicate during this process and the full array of genes required remains incomplete. To study this process in *Drosophila*, we evaluated glial-mediated synapse elimination during metamorphosis, a developmental period in which the nervous system undergoes substantial remodeling, leading to the elimination of a large number of synapses (33). We evaluated Brp-GFP labeled synapses at head eversion (HE), approximately 12 hours into metamorphosis, when glia have already cleared a substantial number of existing synapses in wild type flies (Fig 5A, B). For these experiments, we modified the ELISA assay to only detect full-length Brp-GFP by substituting the GFP detection antibody for a Brp-directed antibody. This was done because of concern that during synaptic engulfment by glia, Brp-GFP might initially be only partially broken down and GFP signal may persist even after synapses were engulfed. This protocol modification can, in principle, be tailored to any protein of interest with antibodies available to evaluate full-length tagged proteins, or even used to compare full length to cleaved GFP tagged proteins to identify regulators of protein processing events.

**Figure 5.**
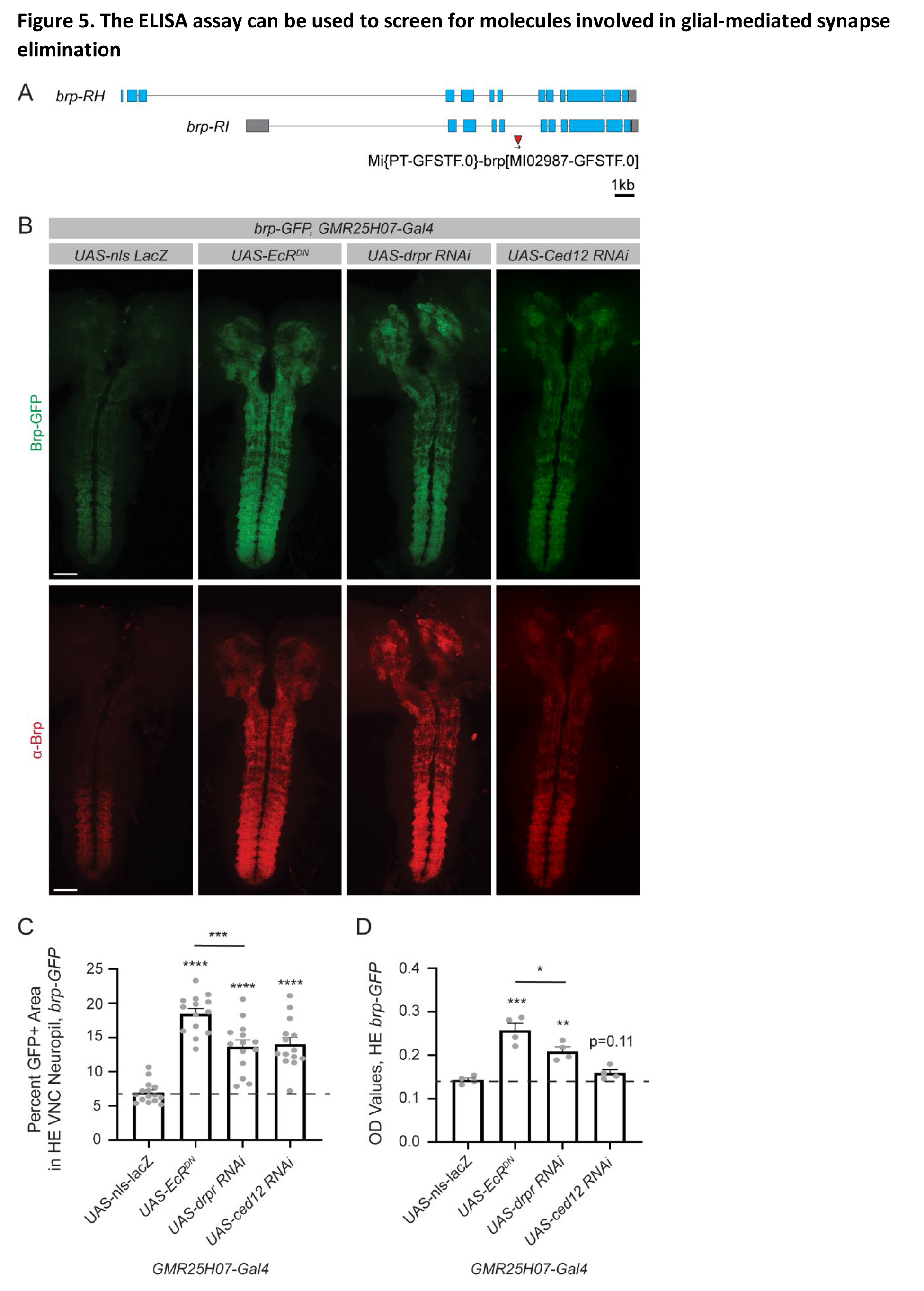
The ELISA assay can be used to screen for molecules involved in glial-mediated synapse elimination. The ELISA assay was used to assess synapse elimination during development using the endogenously tagged synaptic protein, Bruchpilot-GFP (Brp-GFP). Gene expression was manipulated in astrocytes using the *GMR25H07* driver. **(A)** An endogenously tagged Brp protein was previously generated using a MiMIC insertion of GFP into the endogenous *brp* locus. **(B)** The CNS was dissected from pupae at head eversion (HE), approximately 12 hours into metamorphosis. Immunohistochemistry was performed for Brp, and the endogenous Brp-GFP signal was used to evaluate synapses. Images were acquired across the entire CNS and images shown are maximum z projections from 2 stitched 20x images. **(C)** GFP+ area within the VNC neuropil was quantified from single z planes. **(D)** Five dissected CNS were collected for each sample and GFP levels were assessed by ELISA. Graphs represent mean ± SEM and the dotted line indicates the mean of the control *UAS-nls-lacZ* genotype. Statistical comparisons were performed between each experimental genotype and the control. *p<0.05, **p<0.01, ***p<0.001, ****p<0.0001, or as indicated. Scale bar - 50μm.

We first validated that the ELISA assay could detect expected differences in synapses in genotypes known to affect glial-mediated synapse elimination. As expected based on previous work (33), we found that astrocyte expression of the dominant negative EcR^DN^, an RNAi directed against *drpr*, or an RNAi directed against a downstream phagocytic pathway component, *Ced12*, all resulted in increased retention of synapses into metamorphosis by imaging (Fig 5B, C). There were no significant differences evident at larval stages across genotypes (Fig S5B), indicating that these genetic manipulations to astrocytes specifically affected synapse elimination, rather than causing aberrant synaptic development. By ELISA, we were able to detect differences in synapse number at HE with astrocyte expression of EcR^DN^, and *drpr* RNAi, but not with knockdown of *Ced12* (Fig 5D). This indicates that the ELISA can detect physiologically relevant changes in synapse number during development, though it is less sensitive than imaging and quantification to subtle changes in synapse numbers in this context. Similar to what we observed by imaging, no differences in Brp-GFP levels were detected in larval samples by ELISA (Fig S5C). We also validated that quantification of Brp signal by immunohistochemistry showed similar results to those obtained with the endogenously expressed Brp-GFP (Figs S5D, E). Together, these results demonstrate that the ELISA assay can detect expected differences in Brp-GFP levels during glial-mediated synapse elimination in *Drosophila*.

We then used this method to perform a small-scale screen, in order to identify novel genes involved in the process of glial-mediated synapse elimination. Using our ELISA, we identified a class of *syntaxin* genes that all increased synapse retention in metamorphosis to varying degrees (Fig 6A). Syntaxins are known to play roles in vesicular fusion and transport (41) and have been extensively studied in the context of neuronal transmission (42). However, their roles in glia are not well understood. We validated these results by imaging (Fig 6B), which showed similar trends as the ELISA results. Notably, it was difficult to assess the relative differences in synapse numbers among different genotypes by imaging, while they were clearly distinguishable by ELISA. There were no changes in Brp-GFP signal with these RNAi lines at larval stages (Fig S6A, B), indicating that *syntaxins* seem to be specifically important for astrocyte elimination of synapses during metamorphosis.

**Figure 6.**
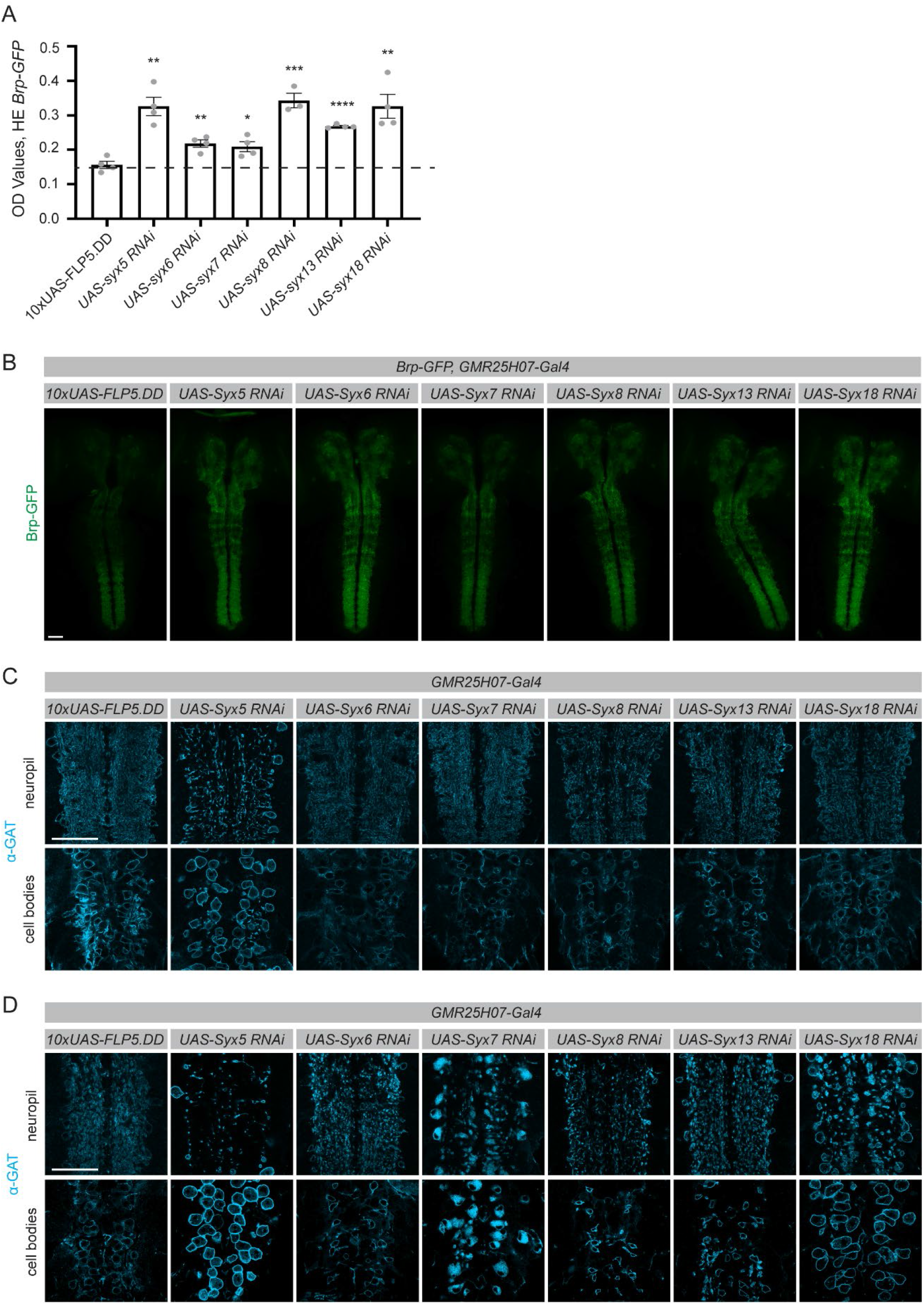
The ELISA assay was used to identify novel genes involved in glial regulation of developmental synapse elimination. A small scale screen was performed to identify novel genes involved in astrocyte-mediated synapse elimination during metamorphosis. **(A)** The CNS was dissected from pupae at head eversion (HE), approximately 12 hours into metamorphosis. Lysates were prepared from 5 CNS for each sample. The ELISA was used to assess endogenously tagged Brp-GFP levels. Genes were knocked down in astrocytes by expressing UAS-driven RNAi lines using the *GMR25H07-Gal4* driver. Statistical comparisons were performed between each experimental genotype and the control *10xUAS-FLP5*.*DD* line, the mean of which is shown as a dotted line. **(B)** Results from the ELISA assay were confirmed by evaluating Brp-GFP in pupae by imaging. Images were acquired at 20x and two images were stitched to show maximum intensity projections through the CNS. **(C)** GAT immunohistochemistry was performed to evaluate the morphology of astrocytes in both the neuropil and cell body. Images were acquired at 63x magnification and a single z plane is shown from a region of the VNC at approximately A1-A3. This was performed in larval samples and **(D)** in pupal samples at 6 hours after puparium formation. Graphs represent mean ± SEM. *p<0.05, **p<0.01, ***p<0.001, ****p<0.0001. Scale bar - 50μm.

To investigate how these Syntaxins might regulate glial-mediated synapse elimination, we examined the morphology of astrocytes during this developmental process. At larval stages, astrocytes infiltrate the neuropil and closely associate with synapses (27). Just 6 hours into metamorphosis (6 APF), astrocytes transform into highly active phagocytes that engulf synaptic material and other cellular debris (33). Interestingly, knockdown of the different *syntaxin* genes had a wide range of effects on this process of astrocytic transformation (Fig 6C,D). Knockdown of *syntaxins 5* and *8* showed deficits in astrocyte morphology that were evident even at larval stages, and continued to show dramatic loss of infiltration of astrocyte processes into the neuropil during metamorphosis. Knockdown of these genes also produced some of the strongest phenotypes in aberrant synapse retention as measured by ELISA. In contrast, *syntaxin 7* knockdown resulted in relatively normal astrocyte morphology at larval stages, but also exhibited reduced process infiltration of the neuropil during metamorphosis and accumulation of the membrane marker GAT in cell bodies. Despite this, synapse elimination was only mildly affected, suggesting that astrocytes do not need to cover the entire neuropil to effectively eliminate most synapses. *Syntaxin 18* knockdown astrocytes also appear relatively normal at larval stages, while in metamorphosis they accumulate large vesicular structures in their processes and exhibit substantially enlarged cell bodies. While astrocytes are clearly are able to undergo morphological transformation during metamorphosis in this condition, the ELISA results demonstrate that they are not effective at phagocytosing synapses. *Syntaxin 6* and *syntaxin 13* knockdown result in relatively normal astrocyte coverage in metamorphosis, but impaired phagocytic function, reflected in a mild defect in synapse elimination. Thus, the class of *syntaxins* identified in this small-scale screen result in diverse changes in astrocyte morphology and function. These data provide proof of principle that the ELISA assay can identify novel molecules involved in the process of glial-mediated synapse elimination, and revealed surprising roles for *syntaxins* in mediating the changes in astrocyte morphology that are required for successful elimination of synapses during *Drosophila* development.

## Discussion

Our work demonstrates that a GFP-based ELISA assay can be used to rapidly perform quantitative screens in an *in vivo* system. In conjunction with the array of tools available in *Drosophila* to express GFP in numerous different subsets of cells, GFP-based transcriptional reporters, and thousands of endogenously GFP tagged proteins, this assay can be used to screen for regulators of an extensive array of cellular and molecular pathways. In this study, we demonstrated the ease of adapting this assay to address a wide variety of questions related to fundamental topics in neuroscience, and were able to validate six completely independent sets of screening paradigms that could be performed with this ELISA platform.

First, we showed that the ELISA assay was capable of detecting GFP expression in a variety of different cell populations, and demonstrated that it could detect changes in GFP signal which correlated with morphological remodeling of *pdf* neurons. One could thus use this paradigm to screen for genes involved in the regulation of circadian neuronal remodeling. In addition, we found that the assay could detect GFP expression in a variety of other cell types, including mushroom body neurons, which undergo plastic changes in response to learning. This assay could potentially be used to screen for genes that modify experience-dependent neuronal plasticity. This assay can also detect signal from neurons in the antennal lobes, which could be used to identify genes involved in the clearance of neuronal debris after injury. These examples represent only a small handful of the subsets of neurons that can be labeled in *Drosophila* using the extensive array of driver lines that have been developed (12-14). This ELISA assay could be used with any drivers that induce high enough levels of expression. Detection of GFP signal within the small number of *pdf* neurons, approximately 150 per brain, suggests that this assay could be used even with drivers that express in small subsets of cells.

We further demonstrate that our ELISA assay can be used to identify genes involved in the morphogenesis of astrocytes. While the complex morphology of many cells in the nervous system has long been a topic of interest, the genes that establish these unique morphologies have been difficult to screen for in a systematic manner. It is challenging to quantitatively distinguish different patterns of infiltration of these densely organized cell types *in vivo*, and even if this were possible, the imaging and analysis would be prohibitively time consuming to perform a large-scale screen. To address these challenges, we paired the ELISA assay with expression of a membrane localized GFP. This is not a direct measurement of morphological complexity, and could also reflect changes in cell number or changes in protein trafficking to the membrane. Nonetheless, in this context, the assay was able to detect changes in GFP signal in response to genetic manipulations known to regulate astrocyte morphology. This could provide a platform for a large-scale screen of genes which are involved in establishing the morphological features of glia. In addition, this same principle could be applied to screen for genes involved in the regulation of other cell types with interesting morphological characteristics which have previously not been possible to study. Here, we used a membrane localized GFP as a proxy to reflect the membrane content of these cells. However, *Drosophila* lines are available with GFP localized to virtually every cellular compartment, and thus the same principle could be used to screen for regulators of mitochondrial density, ER development, lysosomal content, or other subcellular features, in any desired subset of cells.

In addition to evaluating features of cells themselves, we also demonstrate that this assay can be used to assess changes in signaling pathways within desired cellular subsets. Specifically, we showed that the assay could detect predicted changes in GFP in astrocytes expressing a GFP-based transcriptional reporter for activation of the STAT pathway following injury. This could be used to screen for regulators of injury responses in glia, and could be extended to assess glial responses in disease models, which also rely on activation through this pathway. Based on our imaging experiments, this transcriptional reporter produced only a modest amount of GFP in the contexts examined, suggesting that the ELISA could detect relatively weak or subtle changes using other transcriptional reporter systems as well. A wide variety of transcriptional reporters have been developed in *Drosophila* to assess activity of specific transcription factors (20), and could be paired with this ELISA assay to screen to regulators of these diverse transcriptional programs. In addition, transcriptional reporters have been developed that respond to cytoplasmic Ca^2+^ levels, which can be used, among other things, to provide a read out of neuronal activity (17, 18). Thus, ELISAs could be used to screen for modulators of neuronal activity in any desired subset of neurons in response to environmental or genetic perturbations.

Perhaps the largest array of GFP-based tools available in *Drosophila* are endogenously GFP tagged proteins. Thanks to large scale efforts and multiple approaches toward the goal of tagging every conserved protein in *Drosophila* (22-25), there are already thousands of these lines available. We showed that this assay could detect GFP expression in two such lines, expressing Drpr-GFP and Cry-GFP. In both cases we found that the ELISA assay could detect expected changes in the expression of these proteins, even over short time-scales, during the course of development and across circadian time. Genetic knockdown or mutants known to affect expression of these proteins also produced the predicted outcomes by ELISA. These two examples serve as a proof of principle that this assay could be used to evaluate expression of a variety of proteins, and thus could be used to screen for regulators of expression of virtually any protein of interest.

We took advantage of our assay’s capability to detect endogenously tagged proteins to evaluate the contribution of astrocytes to synapse elimination in *Drosophila* by measuring Brp-GFP by ELISA. We evaluated Brp-GFP levels as a proxy to assess synapse number during the process of metamorphosis, a developmental stage when the *Drosophila* nervous system undergoes remodeling and many synapses are eliminated. We first validated that the assay could detect aberrant retention of synapses with knockdown of genes within astrocytes which had previously been shown to regulate this process (33), and indeed we were able to recapitulate these previous findings. We then used this platform to perform a small-scale screen to identify novel regulators of glial-mediated synaptic engulfment, and identified a previously unknown role for *syntaxins* in this process. Upon examining astrocyte morphology, we revealed that these *syntaxin* genes are necessary to regulate the changes in astrocytic phenotypes required to engage in synapse elimination during metamorphosis. Syntaxins 7 and 13 have previously been shown to be associated with vesicles in cultured phagocytes (43), but broader roles for this class of proteins in phagocytic function in the nervous system was unknown. This screening paradigm could be directly extended to evaluate regulators of synapse development, elimination and plasticity in other contexts.

For our analyses of synapses, we used a modified ELISA assay to identify full-length Brp-tagged GFP by pairing GFP coating and Brp detection antibodies. We did this to ensure that we were evaluating intact synaptic material, as there was concern that after engulfment by glia that GFP itself might remain intact for some time within the endosomal pathway. However, this protocol modification could be used in other studies aimed at assessing relative levels of full-length and cleaved versions of different protein products, or even to study regulators of posttranslational protein modifications by pairing GFP with antibodies against phosphate, ubiquitin, glycosides, or other desired functional groups. Together, these examples demonstrate that this GFP-based ELISA method can be used to screen for regulators of a wide variety of specific cellular and molecular pathways in an *in vivo* system.

Our screening method offers several advantages over the current methods that are used to perform screens aimed at identifying genes involved in specific cellular and molecular pathways, which most frequently rely on imaging. First, samples are simply frozen at the time of collection, and thus can be collected over time, and the assay itself later run on desired batches of samples together at the investigator’s convenience. For adult heads, no dissections were required, so whole flies can be collected and their heads simply dissociated by vortexing. This takes little time, even for very large numbers of flies, and does not necessitate the technical skill required to perform dissections and tissue mounting. We did find that larval and pupal nervous systems needed to be dissected to avoid non-specific background signal using this assay. However, we hope in the future to identify the source of protein interfering with signal from whole animals at these developmental stages to further reduce the time required for these experiments. Regardless, after samples are collected, the assay itself requires little active time, making it substantially faster than mounting, imaging and manually evaluating samples in imaging-based screens. While the exact time savings will vary based on the set up of the screen, we anticipate that running an ELISA-based assay for 1500 lines would take weeks, while most imaging based screens require years to complete. In addition to being faster, this ELISA based screening method also requires no equipment beyond a colorimetric plate reader and basic supplies commonly available in the laboratory. The cost of performing a screen using this method is also much lower than an imaging based screen. No microscope time is required, and we estimate that material costs of performing the ELISA assay would not exceed $1500 for 1500 lines. Finally, this screening method produces quantitative results. Thus, one does not have to rely on multiple investigators to employ consistent qualitative and subjective scoring criteria to establish hits from a screen, and one can go back at a later time to assess the precise relative strength of the effects of additional genes or conditions within the original screening dataset. Together, this ELISA-based screening assay offers a rapid, low-cost, accessible and quantitative method that can be easily adapted to address a wide variety of cellular and molecular questions.

It is important to note that this method also has limitations. In imaging-based screens, one can see patterns of multiple different phenotypes. The ELISA simply provides a read out of the concentration of GFP molecules, changes in which may indicate a range of phenotypes depending upon how the screen is designed. For example, when we assayed astrocyte morphology in this study, the ELISA produced nearly identical results in conditions in which *ths* and *pyr* were overexpressed. However, by imaging, it was clear that while *ths* overexpression seemed to promote relatively uniform increases in astrocyte density within the neuropil, *pyr* overexpression resulted in highly concentrated clumps of astrocyte processes, and disruption of the neuropil structure. Thus, we anticipate that possible hits identified in ELISA based screens will need to be validated using imaging or other modalities to extract more information about how genes identified as hits might act. This assay would also not be compatible with most MARCM-based screening methods due to the stochastic nature of recombination in this system, and thus could not study questions that require use of a clonal system in which the number of labeled cells varies from animal to animal. In addition, we found that the assay could not be used to reliably compare differences between different developmental stages of the fly. For example, Brp levels in pupal samples should be strongly reduced relative to larvae, but we detected only modest changes in signal among these samples. So, while results within each stage of the life cycle were internally consistent, we could not compare between these groups. The same may be true for comparisons across different tissue types within the fly as well, as we have not tested this directly. This likely could be rectified in the future by establishing different conditions for lysate preparation, but we were not able to identify those here. Finally, while the ELISA is sufficiently sensitive to detect relatively low levels of GFP, this screening method is not an ideal platform to study things with very low levels of expression in very small populations of cells. Thus, not all cells and proteins will be amenable to screening using this technique.

We have demonstrated the utility of a GFP-based ELISA for screening in the nervous system of *Drosophila*, but this method could also be extended to use in other applications. Certainly, there is no reason that this method should work only in the nervous system, and it should be broadly applicable for use in other tissues. While we have used *Drosophila* to demonstrate its utility, this assay could also be adapted to work in other organisms that are amenable to forward genetic screens, such as *C. elegans*. In this paper, we developed the ELISA to work with GFP, since numerous GFP-based tools are already available, such that this assay could be used with virtually no troubleshooting of the assay itself for a wide variety of applications. However, ELISAs can work with many antibodies, and in cases where two antibodies are available against a given protein that recognize different epitopes, an ELISA assay may be able to be optimized to detect that native protein. In this case, there would be no need to use any lines that express GFP tags, and the protein itself could be measured directly. This approach and the GFP-based ELISA developed in this work could also be used for purposes beyond screening. These assays produce highly quantitative readouts of protein concentrations, and thus can be used as a faster, more quantitative and more sensitive tool to quantify relative changes in protein levels compared to Western blots. This could be used to assess efficiency of knockdowns at the protein level if a GFP-tagged version of that protein is available, assess the relative strength of different driver lines by expressing GFP under their control, or simply to assess the effect of targeted manipulations on the levels of a given protein of interest.

In this study, we have demonstrated that a GFP-based ELISA can be used to rapidly screen for regulators of a wide variety of cellular and molecular pathways in *Drosophila*, demonstrating its utility as a tool to perform *in vivo* forward genetic screens. This assay makes *Drosophila* an even more powerful system to perform forward genetic screens that aim to understand a wide variety of targeted phenotypes and processes. This method promises to make these screens faster to validate and perform, accelerating the discovery of novel genes involved in diverse biological processes.

## Materials and Methods

### Fly Strains and Maintenance

Flies were maintained at 25°C on cornmeal molasses media. Flies were kept in an incubator with 12 hour light-dark cycle. However, no additional precautions were used to ensure strict circadian entrainment. The flies used in this study are as follows: *pUAST-mCD8::GFP(44), elaV-Gal4* (45), *Tab[201Y]-Gal4* (46), *Orco-Gal4* (47), *pdf-Gal4* (48), *repo-Gal4*(49), *Alrm-Gal4* (50), *GMR25H07-Gal4* (12), *w*^*1118*^, *UAS-nls-lacZ* (BL3955), *10xUAS-FLP5*.*DD* (51), *UAS-λ-htl* (52), *UAS-ths* (53), *UAS-pyr* (54), *UAS-EcR*^*F645A*^ (55), *per*^*01*^ (56), *10xSTAT92E-dGFP* (19), *Drpr-GFP* (57), *Cry-GFP* (34), *UAS-htl-RNAi* (VDRC #GD27180), *UAS-pWiz-drpr RNAi7b*(58), *UAS-Ced12 RNAi* (VDRC # 107590), *UAS-syx5-RNAi* (VDRC # KK100952), *UAS-syx6-RNAi* (VDRC # 109340), *UAS-syx7-RNAi* (VDRC # KK101990), *UAS-syx8-RNAi* (VDRC # KK101612), *UAS-syx13-RNAi* (VDRC # KK111650) and *UAS-syx18-RNAi* (VDRC # KK101345).

### ELISA Assay

#### ELISA Sample Preparation

For larval and pupal samples, the nervous system was dissected in PBS containing 0.1% Tween-20 (0.1% PBST) and kept in 1.5mL Ependorf tubes on ice until all samples were collected. Samples were frozen and stored at -80°C until use.

For adult samples, flies were anesthetized using CO2 and collected in 1.5mL Ependorf tubes. Flies were then frozen on dry ice and stored at -80°C. To remove heads, tubes were vortexed at high speed for 15-20 seconds. Tubes were then inverted onto a white index card and the desired number of heads for each sample collected in fresh tubes. These tubes were then stored at -80°C until use. We detected no substantial changes in signal in any of the applications used in this study when heads were thawed at this collection step. If large numbers of heads are required, flies can be collected in 15mL conical tubes, frozen, vortexed and passed through a series of two sieves (size no. 25 and no. 40 respectively). Heads will settle into the second sieve and can be gently removed using a paintbrush.

At the time of the assay, all samples were removed from -80°C and placed on ice. 120μl of 0.1% PBST was added to each sample and samples were homogenized using a pestle with a hand-held homogenizer in 1.5mL Ependorf tubes for 5 seconds. Longer homogenization times, addition of protease inhibitors and sonication were tested, but had no impact on signal in the applications used in this study. Samples were placed on ice until all samples had been processed. Samples were centrifuged at 13,000 rpm for 10 min at 4°C and placed on ice until loading into the plate. See summary in Fig 7A.

**Figure 7.**
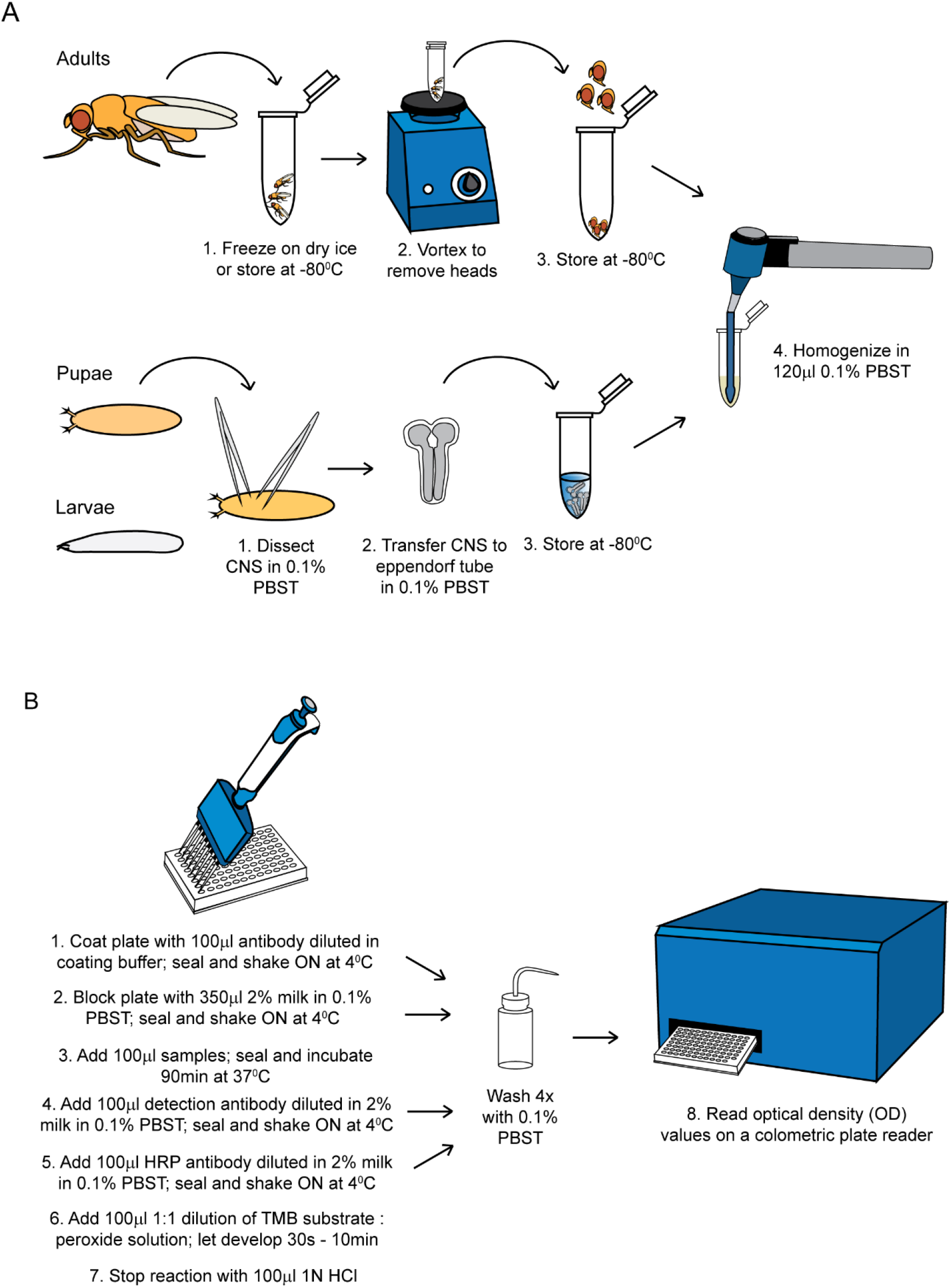
An overview of the sample preparation and ELISA assay protocols. **(A)** An overview of the procedure for preparing of *Drosophila* tissue for use in the ELISA assay is shown. In this study, adult fly heads were processed whole, while the CNS from larvae and pupae were dissected. Samples can be stored for at least several weeks at -80°C before homogenates are prepared and loaded into ELISA plates. **(B)** An overview of the protocol for the ELISA assay is shown. If all samples are run in triplicate, each 96 well plate can accommodate 30 experimental samples, a blank sample (0.1% PBST) and a control to standardize OD values across multiple plates. This control can be a standard concentration of recombinant GFP or a biological control sample that is run on all plates. While this should be sufficient to compare relative values across plates for a screen, a standard curve of recombinant GFP can also be run on each plate in order to obtain a readout of the concentration of GFP molecules for each sample.

#### ELISA Assay

Plates (Immunolon 2HB flat bottom, Thermo #3455) were coated in 100μl buffer (30mM sodium carbonate, 70mM sodium bicarbonate in ddH_2_O, pH 9.6) containing a 1:1000 dilution of chicken αGFP antibody (abcam #ab13970). Our results in Figs 1A, B indicate that lower concentrations, down to 1:5000 could also be used, but because of the wide range of applications and signal intensities anticipated throughout our studies, we used an excess of antibody to ensure we captured even samples with relatively high signal within the assay’s linear dynamic range. We also tested alternative antibodies and found that a chicken αGFP antibody (Aves #1010) produced slightly lower signal with a 1:5000 antibody dilution, but is a lower cost alternative that would also be suitable for most applications. The plate was sealed using Glad Press-n-Seal and kept on an orbital shaker at 4°C overnight. At all steps, we found that the assay was tolerant of a variety of shaking speeds, but typically used a setting of approximately 40 rpm. The plate cover was removed and the plate washed with 0.1% PBST. For washes, the plate was inverted over a sink and tapped on a paper towel to remove most of the liquid from the wells. The wash solution was applied using a squirt bottle aimed at the sides of the wells. The first and last wash were performed for 4 minutes on an orbital shaker at room temperature (RT) and two additional quick washes were performed in between in which wash solution was added and then removed from the plate immediately. We did find that substantially extending wash times resulted in reduced signal. After these washes, 350μl of 2% non-fat dry milk (Carnation) in 0.1% PBST was added to each well. The plate was sealed and placed on an orbital shaker at 4°C overnight. The seal and blocking solution were removed and the plate washed as described above. After the last wash solution was removed from the plate, samples were added. In our validation experiments, we found that 0.1% PBST alone produced a signal indistinguishable from that of lysates from flies that did not express GFP, and thus used 0.1% PBST was used as our negative control sample. Recombinant GFP (abcam #ab84191) was diluted in 0.1%PBST. Samples should be loaded quickly to prevent the plate from drying out, but we did perform optimization experiments in which samples were not loaded for up to 45 minutes and detected no substantial differences in signal. The plate was then sealed and placed in an incubator at 37°C for 90 minutes. The plate was then inverted to remove the sample lysates, and washed as described above. 100μl of 2% milk prepared in 0.1% PBST containing 1:1000 concentration of mouse αGFP antibody (Life Technologies #A-11120, reconstituted as instructed, then diluted 1:1 in glycerol to a final concentration of 0.1 mg / mL and stored at -20°C) was added, the plate sealed and placed on an orbital shaker at 4°C overnight. This solution was then removed and the plate washed as described above. A 1:2000 solution of donkey α mouse HRP (Jackson ImmunoResearch #715-035-150) was then added in 2% milk in 0.1% PBST, the plate sealed and placed on an orbital shaker at 4°C overnight. This solution was then discarded, the plate washed and 100μl of a 1:1 solution of TMB Solution : Peroxide Solution (TMB substrate Kit, Thermo #34021) added. The assay was allowed to develop for 30 seconds – 10 minutes, depending on the intensity of the signal for each given application. After this time, 100μl of 1N HCl was added to each well to stop the reaction and the plate briefly tapped to mix. The plate was then read using a colorimetric plate reader (ClarioStar Plus, BMG LabTech) at 450 and 630nm. See summary in Fig 7B. In Figures 5 and 6, a similar protocol was used, except that 5% milk was used in place of 2% milk throughout the protocol and the mouse αGFP detection antibody was substituted with a 1:250 concentration of mouse αbrp (nc82, DSHB).

#### ELISA Analysis

The signal for each well was determined by subtracting the 630nm reading from the 405nm reading. The average of the blank wells in each plate was then determined and subtracted from all the samples. These numbers represent the optical density (OD) values reported for each sample throughout the manuscript. In cases where experiments from multiple plates were combined (Fig 1G, Fig 4C), samples from the second experiment were scaled, either by comparing to a GFP standard curve run on the same plate (Fig 1G) or by determining the scale using a set of common internal control samples (Fig 4C) to scale to OD values so they could be compared across experiments.

#### ELISA Kit

In Fig S1C, a GFP ELISA kit was used (abcam #ab171581). This assay was performed following the manufacturer’s instructions, with the exception of sample preparation. Samples were prepared by homogenizing in PBS with 0.1% Tween-20 as described above.

### Immunohistochemistry

Third instar larvae were collected and placed in PBS containing 4% PFA on ice, and then fixed for 20 min while nutating at RT for the experiments shown in Fig.S5 and S6. In the experiments in Fig.2 and S2, larvae were collected in 0.1% Triton X-100 (0.1% PBST) on ice. The CNS was dissected and kept in 0.1% PBST for up to 20 minutes. Samples were transferred to 4% PFA in PBS and fixed for 20 minutes while nutating at RT. In all experiments, after fixation, samples were washed 2 times in 0.1% PBST and transferred to 0.1% PBST containing primary antibodies and incubated for 2 days at 4°C on a shaker. Washes were performed 3 times for 5 minutes at RT with 0.1% PBST and samples were then placed on a shaker in 0.1% PBST overnight at 4°C. Samples were transferred to 0.1% PBST containing secondary antibodies overnight at 4°C, washed and stored in 0.1% PBST overnight or up to 3 nights at 4°C. Samples were mounted in Vectashield antifade reagent (Vector Laboratories). The following primary antibodies were used when staining larvae: α-Brp (nc82 1:50, Developmental Studies Hybridoma Bank), α-GFP (abcam 1:1000), α-GAT (1:3000)(27).

For samples collected at 2 and 6 hours after puparium formation (APF), white prepupae were collected and maintained on a 1% agarose plate in a 25°C incubator for the appropriate time and dissected. For samples collected at head eversion (HE), prepupae were placed on the 1% agarose plate. Every 10 minutes the CNS of animals that head everted were dissected. All dissected samples were placed in PBS with 0.1% Triton X-100 (0.1% PBST) on ice. After this time, samples were prepared as described for larval samples. The following primary antibodies were used for staining pupae: α-Brp (nc82 1:50, Developmental Studies Hybridoma Bank), α-GAT (1:2000) (27), α-Draper (1:400, Developmental Studies Hybridoma Bank) Adult brains were collected from flies at 10-12 days after eclosion, dissected and stained as previously described (12). The cuticle of each head was opened and brains removed in PBS with 0.5% Triton X-100 (0.5% PBST). After dissection, brains were collected in 0.5% PBST on ice. Within 20 minutes, brains were moved into 1.2% PFA in PBS and were fixed for 24 hours at 4°C on a nutator. Brains were washed 3 times in 0.5% PBST for 15 minutes at RT. Brains were transferred into 0.5% PBST with primary antibody and nutated at 4°C for 3-5 days. Brains were washed and transferred to 0.5% PBST containing secondary antibodies and nutated at 4°C for 3-5 days, washed and mounted in Vectashield antifade reagent (Vector Laboratories). The following primary antibodies were used for staining of adult brains: α-Brp (nc82 1:30, Developmental Studies Hybridoma Bank), α-GFP (abcam 1:1000), α-GAT (1:3000) (27).

### Imaging

Images were acquired on a Zeiss 880 confocal system. The magnification, conditions and processing for each image type is indicated in the figure legends. Images were stitched, where indicated, using Zen Software.

### Image Analyses

The investigator performing the image analysis was blinded to genotype and condition. Z stacks were opened in Image J and a single slice was chosen for each sample which contained neuropil across the length of the VNC. The investigator outlined the neuropil, and the total area was measured. Each image was manually thresholded and the total positive area recorded. The percent area was calculated from these measures and were used for analysis.

### Statistical Analysis

Statistics were performed using GraphPad Prism software. Significant outliers were identified using a Grubb’s test with an α of 0.05 and were omitted from further analysis. All other data points are plotted on each graph. Analyses between two samples were compared using unpaired, two-sided t-tests. Equal variances and Gaussian distributions were assumed. Correlation analyses were analyzed using a simple linear regression. Data are plotted as mean ± SEM and values of p<0.05 were considered significant.

## Acknowledgements and Funding Information

The authors would like to thank members of the Freeman lab for their valuable feedback on this work. This work was supported by NIH grant R37-NS053538 to MRF, NIH grant RO1-NS112215 to MRF, NIH grant F32-NS117647 to TRJ and by Damon Runyon Fellowship 2329-18 to YK.

## Figures and Figure Legends

**Figure S1.**
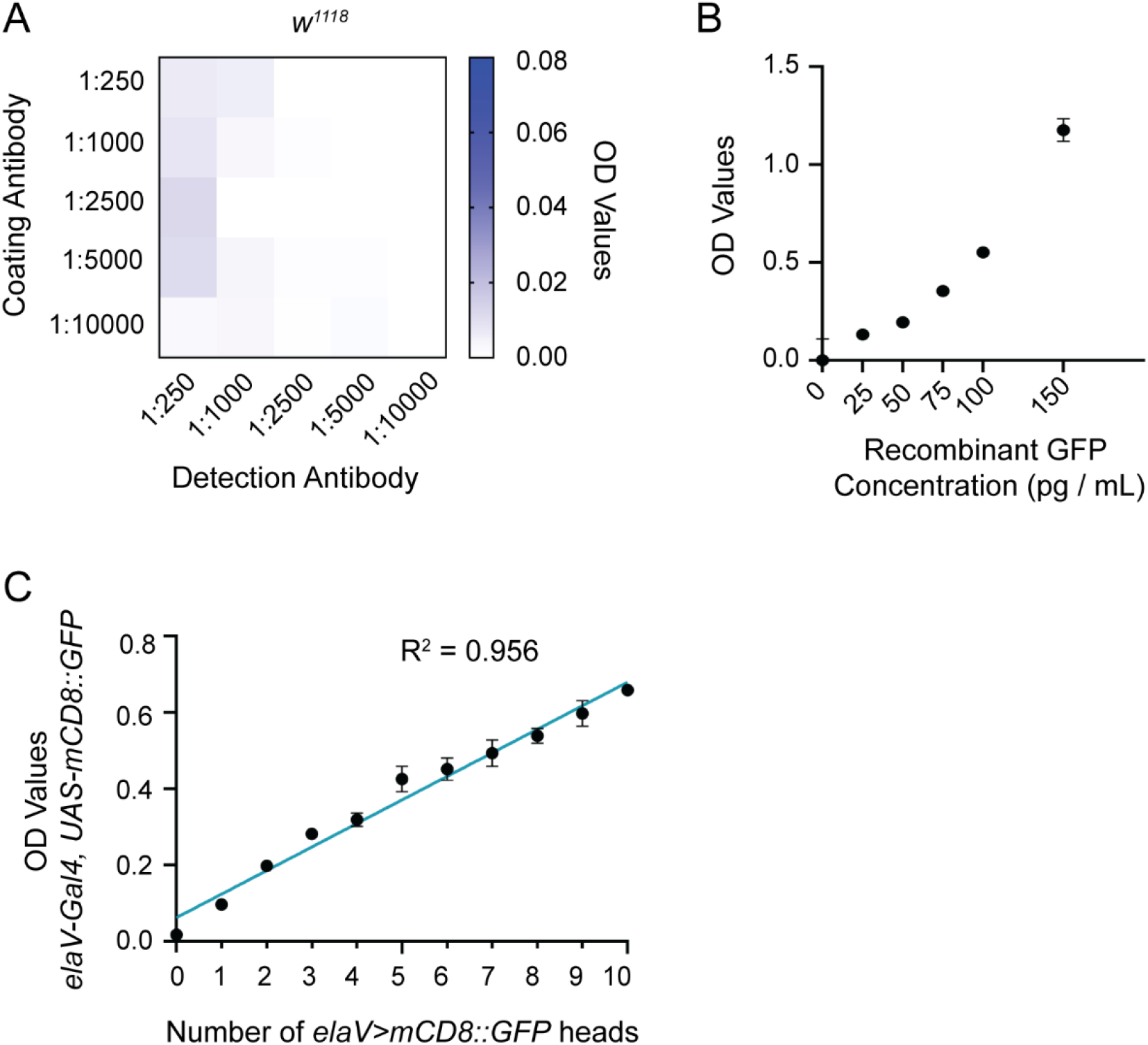
**(A)** The same data shown in Fig 1B are represented, but the scale of the OD values is decreased by a factor of 10 to show the relative background signal in these different antibody conditions. **(B)** OD values were recorded from recombinant GFP after the assay was allowed to fully develop for 10 minutes. **(C)** A commercially available ELISA kit was used to measure signal from lysates prepared from different numbers of *elav>mCD8::GFP* expressing flies, indicated on the x-axis. As in Fig 1D, 10 heads were prepared for each sample with the remaining number of heads from *w*^*1118*^ flies. The correlation analysis was performed using a simple linear regression.

**Figure S2.**
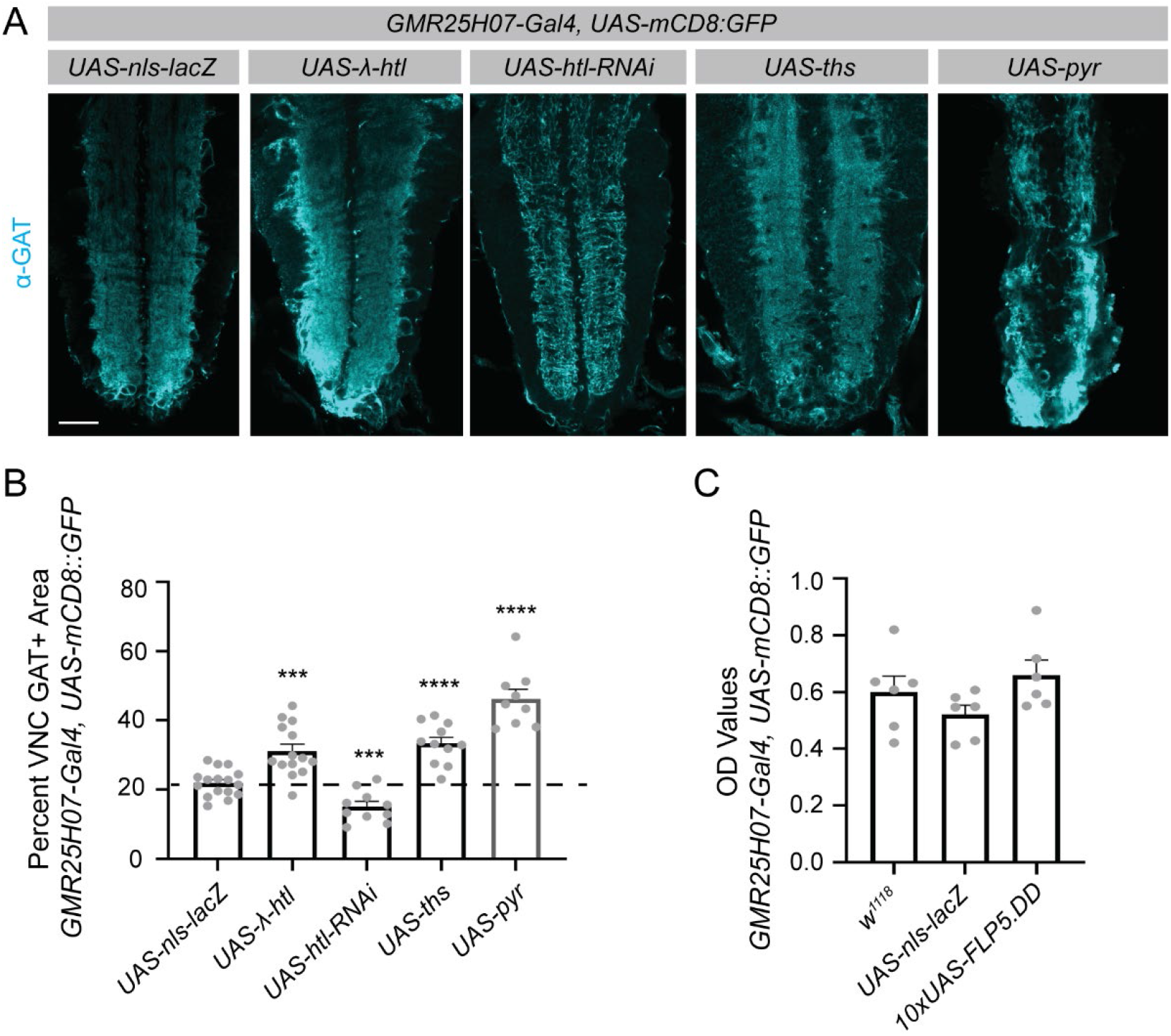
**(A)** The samples shown in Fig 2A were also stained using an antibody against the astrocyte membrane marker GAT. Images represent single z planes through the larval VNC. **(B)** GAT+ immunoreactive area within the VNC neuropil was quantified as above for each genotype. Each experimental genotype was compared to the *UAS-nls-lacZ* control (mean depicted as a dotted line). **(C)** ELISA analyses were performed on lysates prepared from 3 dissected larval CNS for each sample. In addition to driving membrane localized GFP, the *GMR25H07* driver also drove expression of 3 control samples expressing varying numbers of UAS sequences: *w*^*1118*^ (no UAS), *UAS-nls-lacZ* (5xUAS) and *10xUAS-FLP5*.*DD* (10xUAS). There were no statistical differences detected among these groups. Graphs represent mean ± SEM. ***p<0.001, ****p<0.0001. Scale bar - 50μm.

**Figure S5.**
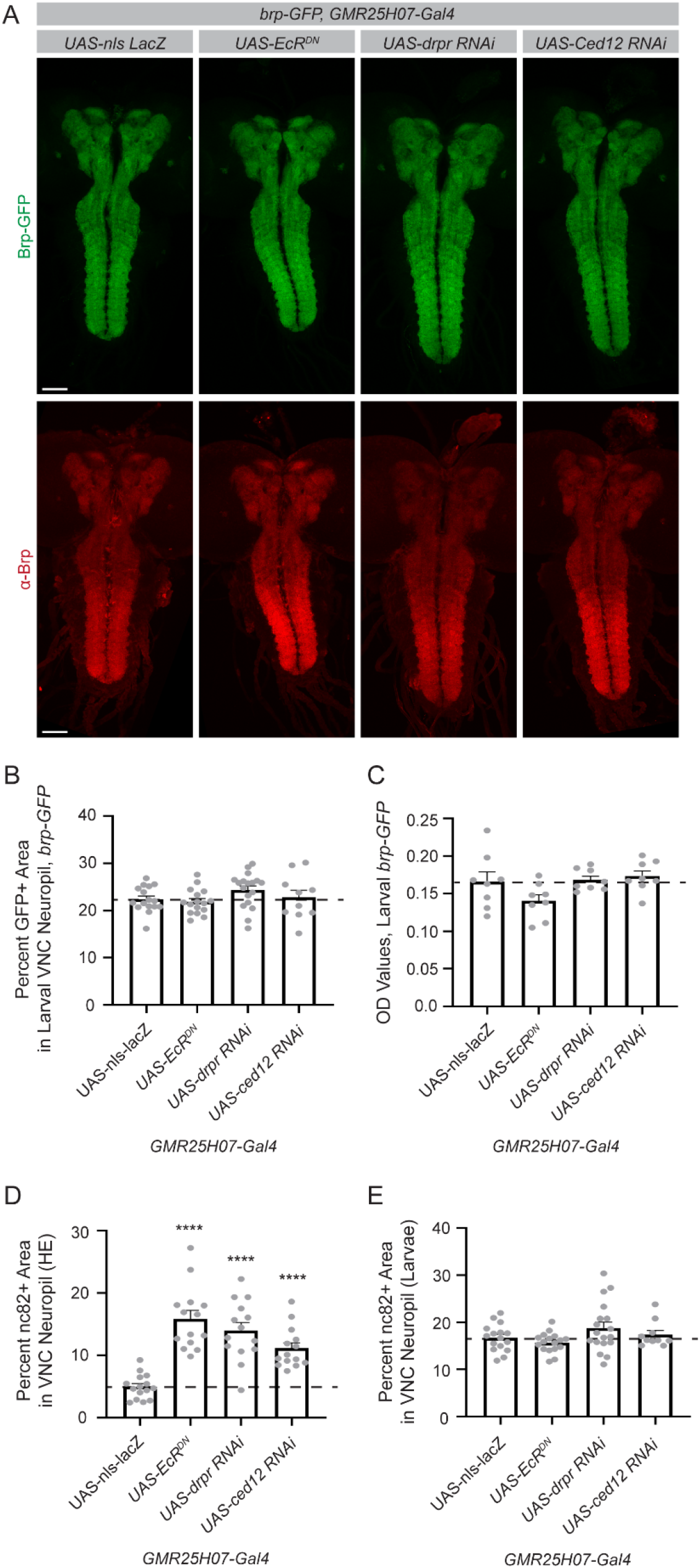
**(A)** Brp-GFP was ubiquitously expressed and gene expression in astrocytes manipulated using the *GMR25H07* driver. The CNS was dissected from larvae and stained for Brp. The GFP represents the endogenous Brp-GFP signal. Images were acquired at 20x and two images were stitched to show a maximum intensity projection through the CNS. **(B)** GFP+ area within the larval VNC neuropil was quantified from single z planes. **(C)** Five dissected larval CNS were prepared for each sample. GFP levels in these lysates were assessed by ELISA. **(D)** The percent area of the VNC neuropil occupied by Brp, as measured by immunohistochemistry using the α-Brp antibody nc82, was quantified in pupal samples at HE and **(E)** in larval samples, as described for GFP area in B. Graphs represent mean ± SEM and the dotted line indicates the mean of the control *UAS-nls-lacZ* genotype. Statistical comparisons were performed between each experimental genotype and this control sample. *p<0.05, **p<0.01, ***p<0.001, ****p<0.0001, or as indicated. Scale bar - 50μm.

**Figure S6.**
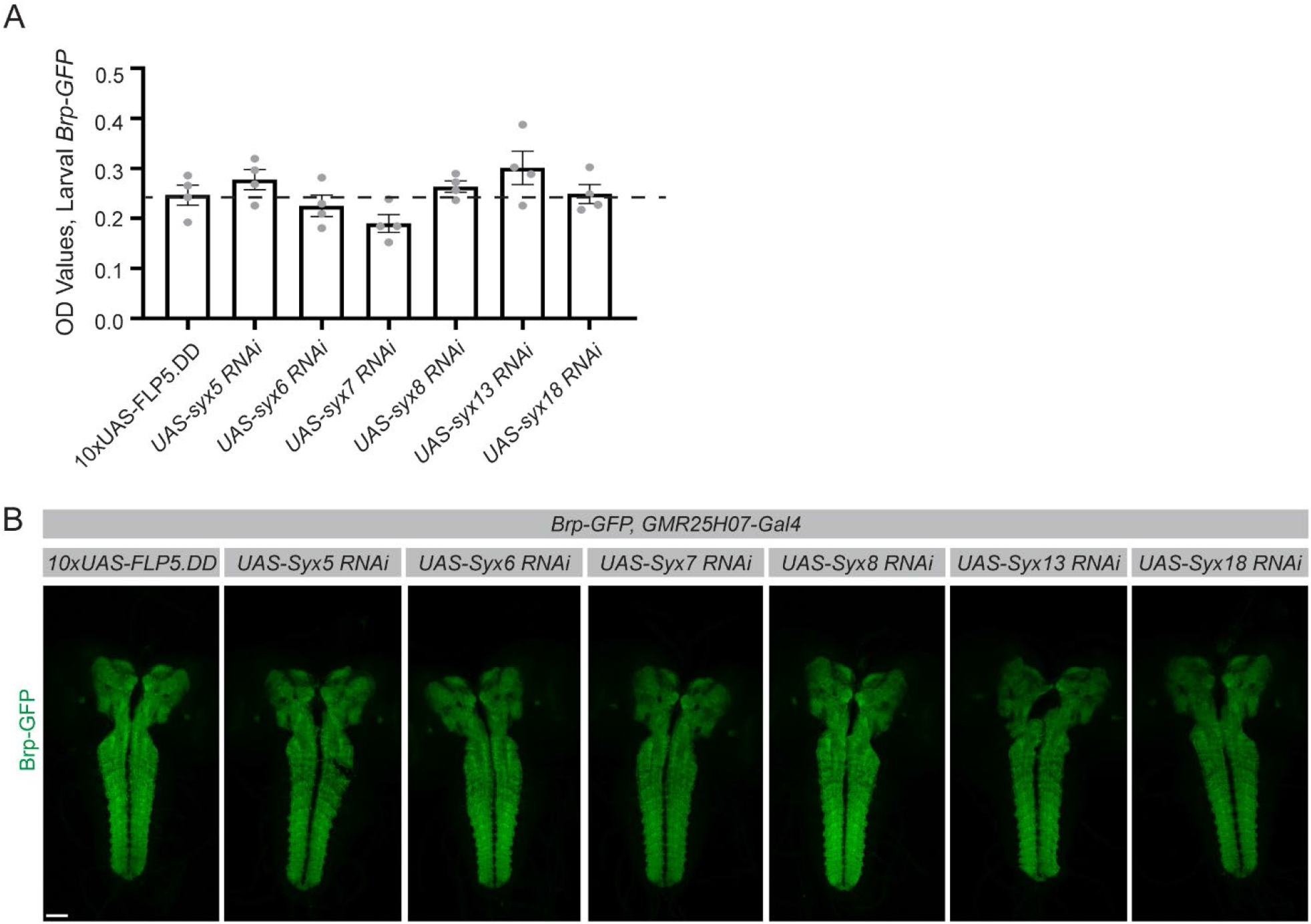
**(A)** Five larval CNS were dissected for each sample. Brp-GFP signal was evaluated by ELISA. The dotted line represents the mean of the control *10XUAS-FLP5*.*DD* genotype and statistics were performed between each experimental genotype and this control sample. There were no statistically significant differences identified among any of the genotypes. **(B)** Larval CNS were dissected and Brp-GFP evaluated by imaging. Images are maximum intensity z projections and 2, 20x images were stitched to show a maximum intensity projection of a z stack through the entire CNS. Graphs represent mean ± SEM. Scale bar - 50μm.

## References

1. D. St Johnston, The art and design of genetic screens: Drosophila melanogaster. Nature Reviews Genetics 3, 176–188 (2002).

2. P. Ligoxygakis, N. Pelte, J. A. Hoffmann, J.-M. Reichhart, Activation of Drosophila Toll during fungal infection by a blood serine protease. Science 297, 114–116 (2002).

3. M. Gans, C. Audit, M. Masson, Isolation and characterization of sex-linked female-sterile mutants in Drosophila melanogaster. Genetics 81, 683–704 (1975).

4. S. Benzer, Behavioral mutants of Drosophila isolated by countercurrent distribution. Proceedings of the National Academy of Sciences of the United states of America 58, 1112 (1967).

5. Y. Dudai, Y.-N. Jan, D. Byers, W. G. Quinn, S. Benzer, dunce, a mutant of Drosophila deficient in learning. Proceedings of the National Academy of Sciences 73, 1684–1688 (1976).

6. R. J. Konopka, S. Benzer, Clock Mutants of <em>Drosophila melanogaster</em>. Proceedings of the National Academy of Sciences 68, 2112–2116 (1971).

7. C. Nüsslein-Volhard, E. Wieschaus, Mutations affecting segment number and polarity in Drosophila. Nature 287, 795–801 (1980).

8. M. A. Simon, D. D. L. Bowtell, G. S. Dodson, T. R. Laverty, G. M. Rubin, Ras1 and a putative guanine nucleotide exchange factor perform crucial steps in signaling by the sevenless protein tyrosine kinase. Cell 67, 701–716 (1991).

9. F. B. Gao, J. E. Brenman, L. Y. Jan, Y. N. Jan, Genes regulating dendritic outgrowth, branching, and routing in Drosophila. Genes Dev 13, 2549–2561 (1999).

10. M. Seeger, G. Tear, D. Ferres-Marco, C. S. Goodman, Mutations affecting growth cone guidance in drosophila: Genes necessary for guidance toward or away from the midline. Neuron 10, 409–426 (1993).

11. L. J. Neukomm, T. C. Burdett, M. A. Gonzalez, S. Züchner, M. R. Freeman, Rapid in vivo forward genetic approach for identifying axon death genes in <em>Drosophila</em>. Proceedings of the National Academy of Sciences 111, 9965–9970 (2014).

12. A. Jenett et al., A GAL4-driver line resource for Drosophila neurobiology. Cell reports 2, 991–1001 (2012).

13. B. D. Pfeiffer et al., Tools for neuroanatomy and neurogenetics in Drosophila. Proceedings of the National Academy of Sciences 105, 9715–9720 (2008).

14. B. D. Pfeiffer et al., Refinement of tools for targeted gene expression in Drosophila. Genetics 186, 735–755 (2010).

15. P. T. Lee et al., A gene-specific T2A-GAL4 library for Drosophila. Elife 7 (2018).

16. H. J. Bellen et al., The Drosophila gene disruption project: progress using transposons with distinctive site specificities. Genetics 188, 731–743 (2011).

17. K. Masuyama, Y. Zhang, Y. Rao, J. W. Wang, Mapping neural circuits with activity-dependent nuclear import of a transcription factor. J Neurogenet 26, 89–102 (2012).

18. X. J. Gao et al., A transcriptional reporter of intracellular Ca(2+) in Drosophila. Nat Neurosci 18, 917–925 (2015).

19. E. A. Bach et al., GFP reporters detect the activation of the Drosophila JAK/STAT pathway in vivo. Gene Expression Patterns 7, 323–331 (2007).

20. N. Chatterjee, D. Bohmann, A versatile ΦC31 based reporter system for measuring AP-1 and Nrf2 signaling in Drosophila and in tissue culture. PLoS One 7, e34063 (2012).

21. A. C. Spradling et al., The Berkeley Drosophila Genome Project gene disruption project: Single P-element insertions mutating 25% of vital Drosophila genes. Genetics 153, 135–177 (1999).

22. X. Morin, R. Daneman, M. Zavortink, W. Chia, A protein trap strategy to detect GFP-tagged proteins expressed from their endogenous loci in Drosophila. Proceedings of the National Academy of Sciences 98, 15050–15055 (2001).

23. H. J. Bellen et al., The BDGP gene disruption project: single transposon insertions associated with 40% of Drosophila genes. Genetics 167, 761–781 (2004).

24. M. Buszczak et al., The carnegie protein trap library: a versatile tool for Drosophila developmental studies. Genetics 175, 1505–1531 (2007).

25. K. J. T. Venken et al., MiMIC: a highly versatile transposon insertion resource for engineering Drosophila melanogaster genes. Nature Methods 8, 737–743 (2011).

26. M. P. Fernández, J. Berni, M. F. Ceriani, Circadian remodeling of neuronal circuits involved in rhythmic behavior. PLoS Biol 6, e69 (2008).

27. T. Stork, A. Sheehan, Ozge E. Tasdemir-Yilmaz, Marc R. Freeman, Neuron-Glia Interactions through the Heartless FGF Receptor Signaling Pathway Mediate Morphogenesis of <em>Drosophila</em> Astrocytes. Neuron 83, 388–403 (2014).

28. J. Doherty et al., PI3K signaling and Stat92E converge to modulate glial responsiveness to axonal injury. PLoS Biol 12, e1001985 (2014).

29. R. J. Kelso et al., Flytrap, a database documenting a GFP protein-trap insertion screen in Drosophila melanogaster. Nucleic Acids Res 32, D418–420 (2004).

30. K. J. Venken et al., Versatile P[acman] BAC libraries for transgenesis studies in Drosophila melanogaster. Nat Methods 6, 431–434 (2009).

31. K. J. T. Venken, Y. He, R. A. Hoskins, H. J. Bellen, P[acman]: A BAC Transgenic Platform for Targeted Insertion of Large DNA Fragments in <em>D. melanogaster</em>. Science 314, 1747–1751 (2006).

32. M. Sarov et al., A genome-wide resource for the analysis of protein localisation in Drosophila. Elife 5, e12068 (2016).

33. O. E. Tasdemir-Yilmaz, M. R. Freeman, Astrocytes engage unique molecular programs to engulf pruned neuronal debris from distinct subsets of neurons. Genes Dev 28, 20–33 (2014).

34. P. Agrawal et al., Drosophila CRY Entrains Clocks in Body Tissues to Light and Maintains Passive Membrane Properties in a Non-clock Body Tissue Independent of Light. Curr Biol 27, 2431-2441.e2433 (2017).

35. P. Emery, W. V. So, M. Kaneko, J. C. Hall, M. Rosbash, CRY, a Drosophila clock and light-regulated cryptochrome, is a major contributor to circadian rhythm resetting and photosensitivity. Cell 95, 669–679 (1998).

36. R. J. Kittel et al., Bruchpilot promotes active zone assembly, Ca2+ channel clustering, and vesicle release. Science 312, 1051–1054 (2006).

37. K. Rein, M. Zöckler, M. T. Mader, C. Grübel, M. Heisenberg, The Drosophila standard brain. Curr Biol 12, 227–231 (2002).

38. W. S. Chung et al., Astrocytes mediate synapse elimination through MEGF10 and MERTK pathways. Nature 504, 394–400 (2013).

39. S. Hong et al., Complement and microglia mediate early synapse loss in Alzheimer mouse models. Science 352, 712–716 (2016).

40. D. P. Schafer et al., Microglia sculpt postnatal neural circuits in an activity and complement-dependent manner. Neuron 74, 691–705 (2012).

41. F. Y. Teng, Y. Wang, B. L. Tang, The syntaxins. Genome Biol 2, Reviews3012 (2001).

42. M. K. Bennett, N. Calakos, R. H. Scheller, Syntaxin: a synaptic protein implicated in docking of synaptic vesicles at presynaptic active zones. Science 257, 255–259 (1992).

43. R. F. Collins, A. D. Schreiber, S. Grinstein, W. S. Trimble, Syntaxins 13 and 7 function at distinct steps during phagocytosis. J Immunol 169, 3250–3256 (2002).

44. T. Lee, L. Luo, Mosaic analysis with a repressible cell marker (MARCM) for Drosophila neural development. Trends in neurosciences 24, 251–254 (2001).

45. D. M. Lin, C. S. Goodman, Ectopic and increased expression of Fasciclin II alters motoneuron growth cone guidance. Neuron 13, 507–523 (1994).

46. M. Y. Yang, J. D. Armstrong, I. Vilinsky, N. J. Strausfeld, K. Kaiser, Subdivision of the Drosophila mushroom bodies by enhancer-trap expression patterns. Neuron 15, 45–54 (1995).

47. M. C. Larsson et al., Or83b encodes a broadly expressed odorant receptor essential for Drosophila olfaction. Neuron 43, 703–714 (2004).

48. J. H. Park et al., Differential regulation of circadian pacemaker output by separate clock genes in <em>Drosophila</em>. Proceedings of the National Academy of Sciences 97, 3608–3613 (2000).

49. K. J. Sepp, J. Schulte, V. J. Auld, Peripheral glia direct axon guidance across the CNS/PNS transition zone. Developmental biology 238, 47–63 (2001).

50. J. Doherty, M. A. Logan, O. E. Taşdemir, M. R. Freeman, Ensheathing glia function as phagocytes in the adult Drosophila brain. J Neurosci 29, 4768–4781 (2009).

51. S. Sethi, J. W. Wang, A versatile genetic tool for post-translational control of gene expression in Drosophila melanogaster. Elife 6 (2017).

52. A. M. Michelson, S. Gisselbrecht, E. Buff, J. B. Skeath, Heartbroken is a specific downstream mediator of FGF receptor signalling in Drosophila. Development 125, 4379–4389 (1998).

53. A. Stathopoulos, B. Tam, M. Ronshaugen, M. Frasch, M. Levine, pyramus and thisbe: FGF genes that pattern the mesoderm of Drosophila embryos. Genes Dev 18, 687–699 (2004).

54. S. Kadam, A. McMahon, P. Tzou, A. Stathopoulos, FGF ligands in Drosophila have distinct activities required to support cell migration and differentiation. Development 136, 739–747 (2009).

55. L. Cherbas, X. Hu, I. Zhimulev, E. Belyaeva, P. Cherbas, EcR isoforms in Drosophila: testing tissue-specific requirements by targeted blockade and rescue. Development 130, 271–284 (2003).

56. S. C. Renn, J. H. Park, M. Rosbash, J. C. Hall, P. H. Taghert, A pdf neuropeptide gene mutation and ablation of PDF neurons each cause severe abnormalities of behavioral circadian rhythms in Drosophila. Cell 99, 791–802 (1999).

57. S. Nagarkar-Jaiswal et al., A library of MiMICs allows tagging of genes and reversible, spatial and temporal knockdown of proteins in Drosophila. Elife 4 (2015).

58. J. M. MacDonald et al., The Drosophila cell corpse engulfment receptor Draper mediates glial clearance of severed axons. Neuron 50, 869–881 (2006).

